# Endothelial sphingosine-1-phosphate receptor 1 deficiency exacerbates brain injury and blood brain barrier dysfunction upon subarachnoid hemorrhage in mice

**DOI:** 10.1101/357236

**Authors:** Akira Ito, Hiroki Uchida, Gab Seok Kim, Giuseppe Faraco, Richard Proia, Kuniyasu Niizuma, Teiji Tominaga, Josef Anrather, Costantino Iadecola, Michael J Kluk, Teresa Sanchez

## Abstract

Blood brain barrier (BBB) dysfunction upon ischemia and hypoxia has been implicated in the exacerbation of neuronal injury in stroke. Despite the therapeutic potential of the cerebrovascular endothelium, the limited understanding of the endothelial signaling pathways governing BBB function restricts progress towards developing novel therapeutic approaches specifically targeting the endothelium. Sphingosine-1-phosphate (S1P) is a potent modulator of endothelial function via its receptors (S1PR). Recent human and mouse studies indicate that vasoprotective endothelial S1P signaling via S1PR1 may be impaired in cardiovascular and inflammatory diseases. Herein, we investigated the expression of S1PR1 in the mouse and human cerebrovascular endothelium and the role of endothelial-specific S1PR1 signaling in brain injury in a mouse model of aneurysmal subarachnoid hemorrhage (SAH), the most devastating type of stroke. We found that S1PR1 is the most abundant S1PR transcript in the mouse brain and in mouse and human brain endothelial cells (20-100 mRNA copies per cell). S1PR1 transcripts were significantly enriched (~6 fold) in mouse cortical microvessels compared to total brain. Using the S1PR1-eGFP knock in mouse, we found that S1PR1-eGFP is abundantly expressed in the cerebrovascular endothelium in the mouse brain. A similar pattern of expression was observed in human brain samples. Endothelial specific deletion of S1PR1 in adult mice (*S1pr1 ^flox/flox^xCdh5-Cre^ERT2^*, referred to as *S1pr1^iECKO^*), resulted in exacerbation of brain edema, neuronal injury and worsened neurological outcomes upon SAH compared to *S1pr1* ^flox/flox^ littermates. No differences in the subarachnoid blood, hemostasis or cerebral blood flow changes during and after SAH were found between groups. Mechanistically, *S1pr1^iECKO^* exhibited aggravated BBB dysfunction and increased phosphorylation of myosin light chain (MLC) in isolated cortical microvessels, a downstream effector of the Rho-ROCK pathway implicated in endothelial inflammation and barrier dysfunction. Taken together, our data indicate that S1PR1 is an endogenous protective signaling pathway in the endothelium, critical to maintain BBB function and to mitigate neuronal injury in pathological conditions. Thus, the therapeutic and diagnostic potential of the endothelial sphingosine-1-phosphate pathway in stroke deserves further study.

## INTRODUCTION

Despite decades of intensive research, stroke is still a leading cause of mortality and long-term disability worldwide [1] and novel therapeutic strategies are need. The cerebrovascular endothelium, in coordination with pericytes [2, 3] and astrocytes [4] plays a critical role in the maintenance of the blood brain barrier (BBB). BBB dysfunction has been implicated in the exacerbation of neurovascular ischemic, hypoxic and inflammatory injury in stroke. For instance, the use and effectiveness of current therapies for ischemic stroke (i.e. tissue plasminogen activator, tPA, or mechanical recanalization) are severely limited by the vascular complications of reperfusion injury (i.e. blood brain barrier dysfunction leading to vasogenic edema and hemorrhagic transformation)[5–9]. In hemorrhagic stroke, perihematoma edema and volume expansion is associated with worsened outcomes [10]. In subarachnoid hemorrhage (SAH), blood brain barrier breakdown occurs in the acute phase and correlates with more severe neurological deficits [11–13]. Although the therapeutic potential of the cerebrovascular endothelium is beginning to be recognized [14–17], *the limited understanding of the endothelial signaling pathways governing BBB function/dysfunction in pathological conditions, restricts progress towards developing novel endothelial-targeted therapies*.

S1P, a bioactive lipid very abundant in plasma, is a potent modulator of endothelial and lymphocyte function. Plasma S1P originates from endothelial cells[18, 19] and erythrocytes[20] and it is bound to lipoproteins (e.g. HDL[21, 22] via apolipoprotein M (ApoM)[22], albumin and other chaperones. In mice, S1P signaling via S1PR1 is required for embryonic and postnatal vascular development and maturation in brain and other organs [23, 24] [25]. In adult mice, S1PR1 promotes endothelial barrier function, vascular stabilization and inhibits endothelial inflammation [26–28]. These effects are dependent on the activation of anti-inflammatory signaling pathways (Gi-phosphatidylinositol-3-kinase leading to activation of Rac and Akt) which strengthen endothelial barrier function by promoting actin cytoskeleton dynamics (cortical actin assembly) and adherens junctions assembly to the cytoskeleton. In contrast, S1P binding to S1PR2 induces barrier dysfunction and activates pro-inflammatory signaling pathways (e.g. NFκB) in a Rho-Rho kinase dependent way. S1P-S1PR1 antiinflammatory signaling pathway is predominant in the endothelium when S1P is bound to HDL via ApoM [28] (Reviewed in [17]). Indeed, a good part of the cardiovascular protective effects of HDL depend on its content of S1P [29–31] and ApoM [28], which also has been shown to protect plasma S1P from degradation [22]. Recent studies indicate that S1P anti-inflammatory and vasoprotective signaling via S1PR1 could be limiting in human cardiovascular and inflammatory diseases [31–34] [17]. For instance, in coronary artery disease, the content of ApoM or S1P in the HDL fraction is significantly reduced, and it inversely correlates with the severity of the disease[31, 32]. In addition, in patients suffering inflammatory conditions such diabetes or sepsis, plasma ApoM levels are significantly decreased compared to control patients [35, 36].

Interestingly, S1PR1 also plays a critical role in lymphocyte trafficking and function: mice lacking S1PR1 in hematopoietic cells are lymphopenic due to impaired egress from thymus and lymphoid organs [37]. Indeed, the immunosupressors, Fingolimod (FTY720) [38, 39], and Siponimod (BAF312)[40], which have been approved by the FDA for multiple sclerosis are potent S1PR1 agonists. Given the phenotype of the mice lacking S1PR1 in hematopoietic cells[37] and *in vitro* studies which indicate that FTY720-P induces desensitization of S1PR1 [41, 42], it is accepted that Fingolimod, Siponimod and other agonists [43] and antagonists [44] of S1PR1 lead to immunosuppression via inhibition of S1PR1 signaling in lymphocytes.

Recent studies attempting to understand the role of S1P in cerebral ischemia have relied on the use of these pharmacological modulators of S1PR1 [45, 46], [47] [48], which protected in experimental stroke. Although the mechanisms are not completely clear, FTY720 protection has been attributed to its immunosuppressive effects [48] and its ability to desensitize S1PR1 (reviewed in [17]). *Thus, these studies could not determine the role of endothelial-specific S1P signaling in BBB function modulation and its impact on brain injury and stroke outcomes*. Although immunosuppression is protective in experimental stroke, it is not clear that it could be a good therapeutic strategy in humans, since it also increases the vulnerability of patients to infections [49, 50], a leading cause of mortality in stroke patients[51, 52]. Nevertheless, Fingolimod [53–55] and Siponimod are currently being tested in stroke clinical trials. *Understanding the role of endothelial S1P signaling in stroke will be pivotal for future design of novel vasoprotective therapeutic agents targeting this pathway specifically in the endothelium without compromising the immune response*.

In order to bridge this knowledge gap and given the pathophysiological relevance of the S1P-S1PR1 pathway in humans, in this study we aimed to investigate the expression of S1PR1 in the mouse and human cerebrovascular endothelium as well as the role of endothelial-specific S1PR1 signaling in modulation of blood brain barrier function and its impact on brain injury upon subarachnoid hemorrhage, the most devastating type of stroke. Our results show that S1PR1 is an endogenous protective signaling pathway in the endothelium, critical to maintain BBB function and to mitigate neuronal injury in SAH. They also highlight the critical role of the endothelial S1PR1 pathway in the pathophysiology of cerebral hypoxia-ischemia as well as its therapeutic potential.

## METHODS

### Mice

All animal experiments were approved by the Weill Cornell Institutional Animal Care and Use Committee. Endothelial cell specific *S1pr1* knockout mice (*S1pr1^flox/flox^xCdh5–Cre^ERT2^;* referred to as *S1pr1^iECKO^*) were generated as we have described [56]. *S1pr1^flox/flox^* mice [24] were crossed to *Cdh5–Cre^ERT2^* mice [57] to generate *S1pr1^flox/flox^ Cdh5-Cre^ERT2^* mice. Mice were treated with tamoxifen (Sigma-Aldrich) by oral gavage (75 mg kg^−1^) for 3 days at the age of 8 weeks and used for the experiments 3-4 weeks after tamoxifen treatment. *S1pr1^flox/flox^* littermates treated with tamoxifen were used as control mice. S1pr1-eGFP knock in mice [58] weighing 26-30 g were used for the expression studies. All experiments were performed in male mice.

### Isolation of cortical microvessels

The brain microvessels were isolated as we have previously described [59]. All procedures were performed in a cold room. The brains were collected and rinsed in MCDB131 medium (Thermo Fisher Scientific) with 0.5% fatty acid-free BSA (Millipore Sigma). The leptomeninges, cerebellum, brainstem and white matter were removed on ice. Ipsilateral cortices were homogenized in 8 mL of MCDB131 medium with 0.5% fatty acid-free BSA using a 7-mL loose-fit Dounce tissue grinder (Sigma-Aldrich) with 10 strokes. The homogenates were centrifuged at 2,000 g for 5 min at 4 °C. The pellet was suspended in 15% dextran (molecular weight ~70,000 Da, Sigma-Aldrich) in PBS and centrifuged at 10,000 g for 15 min at 4 °C. The pellet was resuspended in MCDB131 with 0.5% fatty acid-free BSA and centrifuged at 5,000 g for 10 min at 4 °C. The pellet contained the brain microvessels.

### Isolation of mouse primary neurons and mouse mixed glial cells

Mouse (C57BL6) E16.5 embryos were used for primary cortical neuron isolation as we previously described [60]. Cortices were collected and incubated with 0.25% trypsin for 15□min at 37□°C for tissue digestion. Fetal bovine serum was added to stop the trypsin activity. After centrifugation, the supernatant was discarded and DMEM complete media containing 10% FBS and antibiotics were added to the cells. The cell suspension was passed through 70□μm cell strainer (BD Biosciences, Falcon) and was plated on poly-L-lysine (Sigma Aldrich, St Louis, MO) coated dishes with DMEM complete media. The next day, the medium was changed to Neurobasal media containing B27 supplement, antibiotics and GlutaMAX (Life Technologies). At 7 days *in vitro*, the cells (>95% NeuN positive) were harvested for RNA isolation and gene expression quantification by reverse transcription and quantitative PCR (RT-qPCR).

Mixed glial cell culture was prepared using the mouse brains from postnatal day 2 mouse as we previously described [60] Briefly, cerebral cortices were dissected, trypsinized with of 0.25% trypsin–EDTA in Hank’s Balance Salt Solution (HBSS) and were incubated with trypsin (Thermo scientific) and DNase (Worthington) for 15□min at 37□°C. Foetal bovine serum was added to the cell suspension to stop trypsin digestion. Cell suspension was centrifuged and the pellet was resuspended with DMEM containing 20% FBS and antibiotics. Cell suspension was filtered with a 100-μm cell strainer (BD Falcon) into another 50□ml conical tube. Cells were plated onto six-well plates, which were pre-coated with poly-D-lysine. Three days after plating, the media was changed to DMEM containing 10% FBS and antibiotics. Cells were maintained in DMEM containing 10% FBS and antibiotics at 37□°C with 5% CO2, with a medium change every 3 days. At day 10 after plating RNA was isolated for determination of gene expression by RT-qPCR.

### Endovascular perforation SAH surgery

SAH surgery was performed on C57/BL6/J, *S1pr1^iECKO^* mice and their littermate *S1pr1^flox/flox^* controls as we have previously described [61]. In brief, surgery was performed using a dissecting surgical microscope. Temperature was maintained at 36.5–37.5 °C by using a thermostatic blanket (Harvard Apparatus, CMA 450 Animal Temperature Controller) throughout the procedure. Mice were anesthetized with isoflurane inhalation delivered by facemask with O_2_. A 15 mm midline vertical incision was made on the skin in the head. The head was fixed in stereotactic frame and cerebral blood flow was measured by Laser-speckle imager (PeriCam PSI system, Perimed, Sweden). During surgery, mice were in supine position. A 10 mm midline vertical incision was made on the skin in the neck. The common carotid, external carotid and internal arteries were dissected from the adjacent tissue. The superior thyroid artery and the occipital artery were cauterized and cut. The external carotid artery was sutured with a dead knot and cauterized above the suture. A second suture loop was also placed in the external carotid artery just before the bifurcation of the common carotid artery. A slit-knot was placed around the common carotid artery. A small clip was applied to the internal carotid artery and the slip-knot around the common carotid artery was tightened temporally. A small incision was made in the external carotid artery stump. A 5-0 monofilament with a modified tip (0.3 mm × 0.3 mm or 0.3 mm × 1.7 mm) was inserted into the incision and the knot around the external carotid artery was tightened to prevent bleeding. Then, the monofilament was advanced to the common carotid artery, the small clip on the internal carotid artery was removed and the monofilament was guided through the external carotid artery to the internal carotid artery. The knot around the common carotid artery was opened again and the monofilament was introduced to the bifurcation of the internal carotid artery. The monofilament was gently pushed ~1 mm further and then withdrawn to the external carotid artery. The knot around the external carotid artery was loosen and the monofilament was slowly removed. The external carotid artery was quickly ligated to prevent bleeding. The mouse was turned in prone position and induction of subarachnoid hemorrhage was confirmed by reduction of cerebral blood flow by Laser-speckle contrast imager. After the surgery, all animals were maintained in a small animal heated recovery chamber. Two different severities of SAH models were created by changing the tip shapes of a 5-0 monofilament: a rounded tip 0.3mm (width) × 0.3 mm (length) was used for mild SAH model or a tip 0.3 mm (width) × 1.7 mm (length) for severe SAH model. The surgeon and the investigator conducting the analysis were blinded to the genotype of the mice. Animals which did not exhibit a reduction in CBF upon endovascular rupture were excluded from the study.

### RNA isolation, reverse transcription and quantitative PCR analysis (RT-qPCR)

Total RNA from mouse brain, cells and microvessels was prepared using RNeasy Mini Kit (Qiagen, Valencia, CA) as instructed by the manufacturer. To generate cDNA, 100 ng of RNA was reverse transcribed using random primers and SuperScript II RT-polymerase (Invitrogen, Carlsbad, CA). Primers were designed using the Primer Express oligo design program software (Applied Biosystems, Foster City, CA). Real-time quantitative PCR was performed using the SYBR Green I assay on the ABI 7500 Sequence Detection System (Applied Biosystems). PCR reactions for each cDNA sample were performed in duplicate and copy numbers were calculated using standard curves generated from a master template as we previously described. The sequence of the primers used for qPCR are shown in Table 2.

**Table 1.** Physiological variables in *S1pr1^flox/flox^* and *S1pr1^iECKO^* mice.

|  |  |  |  |
| --- | --- | --- | --- |
| Time points | Parameters | *S1pr1<sup>flox/flox</sup>* | *S1pr1<sup>iECKO</sup>* |
| 5 min before SAH | Arterial O<sub>2</sub> saturation (SpO<sub>2</sub>, %) | 98.6 ± 0.2 | 98.4 ± 0.1 |
|  | Heart rate (b.p.m.) | 442.0 ± 23.2 | 459.2 ± 10.7 |
|  | Pulse distention (μm) | 13.8 ± 0.7 | 12.8 ± 0.9 |
|  | Respiratory rate (br.p.m.) | 162.3 ± 4.9 | 155.0 ± 6.7 |
| 5 min after SAH | Arterial O<sub>2</sub> saturation (SpO<sub>2</sub>, %) | 98.6 ± 0.1 | 98.6 ± 0.2 |
|  | Heart rate (b.p.m.) | 539.8 ± 9.9 | 549.7 ± 9.9 |
|  | Pulse distention (μm) | 13.2 ± 1.1 | 12.5 ± 0.9 |
|  | Respiratory rate (br.p.m.) | 133.3 ± 4.4 | 126.8 ± 6.5 |
| bpm, beats per minute; br.p.m., breaths per minute. Pulse distention is a surrogate for pulse pressure. No significant differences were observed in the physiological parameters (arterial O<sub>2</sub> saturation, heart rate, pulse distention and respiratory rate) between *S1pr1<sup>flox/flox</sup>* and *S1pr1<sup>iECKO</sup>* mice, before and after SAH |  |  |  |

**Table 2.**
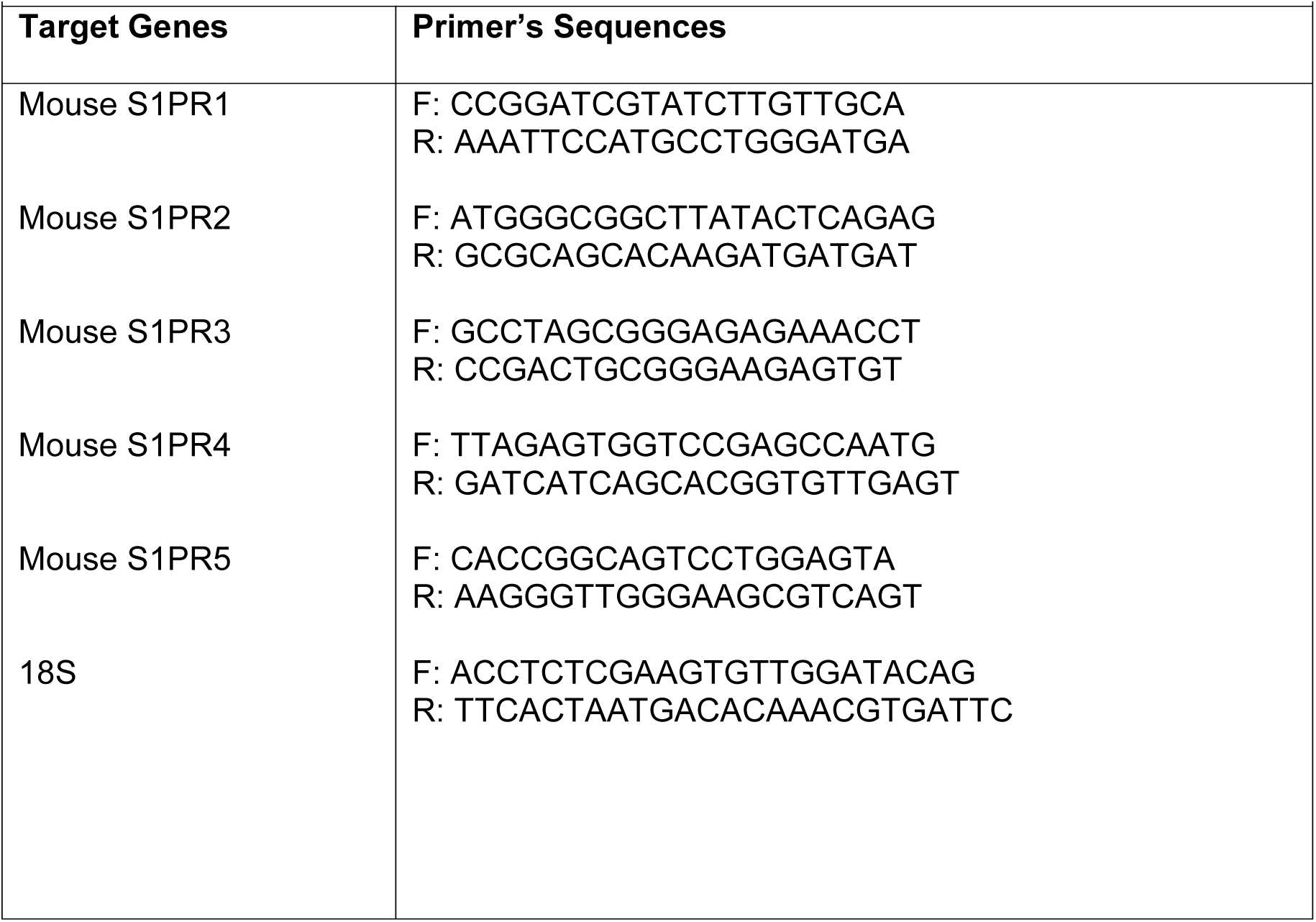
Sequences of the primers used for RT-qPCR

### S1PR1 Immunohistochemistry

Human tissues were retrieved from Brigham and Women’s Department of Pathology archives; this work was approved by the Institutional Review Board (Protocol #2013P001431).

Immunohistochemistry for S1PR1 was performed on an automated stainer (Leica Bond III, Leica Biosystems, Buffalo Grove, IL) using an anti-human S1PR1 rabbit polyclonal antibody (Santa Cruz Biotechnology Inc.) at a final concentration of 1.3 ug/ml. The IHC technique for S1PR1 was validated as we have previously described [62]. 5μm formalin fixed paraffin embedded tissue sections of human frontal cortex were deparaffinized and processed using heat induced epitope retrieval with an EDTA-based buffer (Leica #AR9640) for 20 minutes and incubated with primary antibody for 30 minutes at room temperature. Secondary antibody (polymer) incubation (10 minutes) and diaminobenzidine-based signal generation (10 minutes) were performed per manufacturer’s instructions (Leica # DS9800). Pictures were taken using SPOT Insight Gigabit camera and SPOT Imaging Software (5.1).

### Immunofluorescence staining

Under deep anesthesia, mice were perfused with cold PBS and subsequently with 4% PFA in PBS solution. The brains were removed, postfixed with 4% PFA for 24 h, transferred to 30% sucrose solution in PBS, embedded in OCT media and frozen. Coronal sections were cut (9 µm) in a cryostat. Sections were washed three times with PBS and were then blocked with blocking solution (5 % bovine serum albumin, 0.8 % skim milk, and 0.3 % Triton X-100 in TBS) for 1 h and incubated with the specified primary antibodies in blocking solution overnight on a shaker at 4 °C, followed by the appropriate secondary antibodies and 4’,6-diamidino-2-phenylindole (DAPI) for 1 hours at room temperature and were mounted onto slides. Samples were observed on an FluoView FV10i confocal microscope (Olympus, Japan) (original magnification, × 60).

### Protein extraction from brain microvessels and western blotting

Brain microvascular fragments were lysed in HEPES-RIPA buffer (50 mM HEPES pH 7.5; 1% Triton; 0.5% sodium deoxycholate; 0.1% SDS; 500 mM NaCl; 10 mM MgCl2; 50 mM β-glycerophosphate) with 1x Protease inhibitor cocktail (CalBiochem), 1 mM Na3VO4 and 1 mM NaF and centrifuged at 15,000 r min^−1^ for 15 min. Equal amount of proteins were mixed with SDS sample buffer, boiled and separated on a 4-15% polyacrylamide gel (Bio-Rad), transferred to PVDF membranes (Millipore Sigma), and blocked 5 % milk in 0.1% Tween-20 in TBS. Immunoblot analysis was performed with S1PR1 (1:250; Santa Cruz, cat. no. sc25489), p-MLC (1:1000; Cell Signaling, cat. no. 3671), and β-actin (1:1,000; Santa Cruz, cat. no. sc-1616 HRP) antibodies. Membranes were washed with 0.1% Tween-20 in TBS, incubated with anti-rabbit IgG secondary antibody conjugated to horseradish peroxidase (1:2,000; Cell Signaling), and protein bands were visualized with enhanced chemilumescent (ECL) reagent (Thermo Fisher Scientific) on a Protec OPTIMAX X-Ray Film Processor. Relative band intensities were obtained by densitometric analysis of images using ImageJ software.

### Brain endothelial cell isolation and assessment of deletion efficiency of endothelial *S1pr1 mRNA*

Two weeks after tamoxifen treatment, mice were sacrificed and the brains were collected and rinsed in MCDB131 medium (Thermo Fisher Scientific) with 0.5% fatty acid-free BSA (Millipore Sigma). The cortices were homogenized in MCDB131 medium using a 7-mL loose-fit Dounce tissue grinder (Sigma-Aldrich) with 15 strokes. The homogenate was mixed with same amount of 30% dextran (molecular weight ~70,000 Da, Sigma-Aldrich) in PBS and centrifuged at 4,500 r min^−1^ for 15 min at 4 °C. The pellet was resuspended in MCD131 medium and centrifuged at 2,400 r min^−1^ for 10 min. The pellet was resuspended in Liberase TM solution (3.5 U; Roche) with DNaseI (0.4 U; AppliChem Inc) and digested at 37.0 °C for 90 min. The enzymatic reaction was stopped by adding 2 mM of EDTA and 2% of BSA. After centrifugation at 2,400 r min^−1^ for 10 min, the pellet was incubated in purified Rat Anti-Mouse CD31antibody (MEC13.3) (1 : 100; BD Biosciences) with Dynabeads Sheep Anti-Rat IgG (Invitrogen, cat. no. 11035) for 35 min. CD31 positive endothelial cells were isolated by using DynaMag-2 Magnet (Thermo Fisher Scientific). Total RNA was extracted from the isolated endothelial cells using shredder (Qiagen) and RNeasy Mini Kit (Qiagen) with RNase-free DNase treatment (Qiagen) according to the manufacturer’s instructions. Reverse transcription was carried out using Verso cDNA Synthesis Kit (Thermo Fisher Scientific). Real-time PCR was performed on a real-time PCR system (Applied Biosystems, ABI 7500 Fast) by using PerfeCTa SYBR Green Fast Mix Low ROX. PCR primer sequences for target molecules are in Table 2.

### Grading system for SAH

Blood volume in the subarachnoid space was assessed using the grading system previously reported [63]. Mice were sacrificed under deep anesthesia 24 h after SAH induction and the brains were removed. Pictures of ventral surface of the brain depicting the basal cistern with the circle of Willis and the basilar artery were taken using a stereomicroscope (Olympus, SZX16) equipped with digital camera (Olympus, DP12). The basal cistern was divided into 6 segments and a grade from 0 to 3 was given to each segment: Grade 0, no subarachnoid blood; Grade 1, minimal subarachnoid blood; Grade 2, moderate blood clot with recognizable arteries; Grade 3, blood blot obliterating all arteries. Blood volume was evaluated by a total score ranging from 0 to 18 from six segments.

### Tail bleeding assay

Tail bleeding time was determined as described previously[64]. A mouse was anesthetized with a mixture of ketamine and xylazine, and body weight was measured. The mouse was placed on a heating pad in prone position, the tail tip was about 4 cm blow the body horizon. A distal 5 mm segment of the tail was amputated, and the tail was immediately immersed in PBS pre-warmed at 37 °C. The time to complete arrest of bleeding was determined: complete arrest is no blood flow for 1 minute. The blood volume was determined by hemoglobin assay. Blood cells were separated by centrifuge at 4,000 r/min for 5 min at room temperature, and erythrocytes were resuspended in BD Pharm Lyse (BD Biosciences). After 10 min incubation in the buffer, the lysate was centrifuged at 10,000 rpm/min for 5 min. Hemoglobin concentrations were measured spectrophotometrically at 550 nm using a plate reader (Molecular Devices, SpectraMax M2e).

### Mortality and neurological outcome

Mortality was assessed at 24, 48 and 72 hours after SAH induction. Gross neurological outcome was blindly evaluated before and at 24, 48 and 72 hours after surgery by sensorimotor scoring as described previously [65, 66]. Briefly, a motor score (0 to 12; spontaneous activity, limb symmetry, climbing, and balance) and a sensory score were used. Gross neurological outcome was evaluated by a total score of 4 to 24. Higher scores indicate better neurological outcome.

### Brain water content

Brain edema was determined by using the wet/dry method as previously described [67]. Mice were sacrificed at 72 hours after surgery and the brains were quickly removed and separated into the left and right cerebral hemispheres and weighed (wet weight). The brain specimens were dried in an oven at 55°C for 72 hours and weighed again (dry weight). The percentage of water content was calculated as ([wet weight-dry weight]/wet weight) × 100%.

### Cell death detection

DNA strand breakage during apoptosis after SAH was assessed by phospho-histon H2A.X (Ser 139) immunofluorescence as previously described [68]. Because phosphorylation of histone H2A.X at Ser 139 (γ-H2AX) is abundant, fast, and correlates well with each DNA strand breakage, it is the most sensitive marker that can be used to examine the DNA damage [69, 70]. Mice were deeply anesthetized and perfused with cold PBS and subsequently with 4% PFA in PBS solution. The brains were removed, post-fixed with 4% PFA for 24 h and transferred to 30% sucrose solution. Frozen brains were cut with a 10 µm of thickness by a cryostat (Leica, CM3050 S). The brain slices were blocked with TBS-blocking solution (5% bovine serum albumin and 0.5% Tween 20 in PBS) for 1 hour at room temperature and incubated with phospho-histone H2A.X (Ser 139) (20E3) antibody (1:100; Cell Signaling) in blocking solution overnight on a shaker at 4 °C. Sections were washed three times with 0.5% Tween 20 in PBS and then incubated with goat anti-rabbit IgG Alexa-488 (1:200; Life Technologies, cat. no. A-11008). DAPI (40,6-diamidino-2-phenylindole) staining was used as a counter staining. Sections were imaged by using an FluoView FV10i confocal microscope (Olympus, Japan) (original magnification, × 40). For quantification, the percentage of phospho-histone H2A.X positive cells per DAPI positive cells from three different fields in the ipsilateral cerebral cortex area per mouse (bregma −1.64 to −1.28 mm) were counted and the average values were plotted.

### Assessment of blood brain barrier dysfunction

To assess albumin extravasation, Evans blue dye (EBD) was used because EBD binds to plasma albumin[71]. Mice were anesthetized and 2% EBD (4 ml kg^−1^) was injected in the external jugular vein 21 hours after surgery. After 3 hours circulation, mice were deeply anesthetized and perfused with cold PBS to remove intravascular dye. The ipsilateral hemispheres were removed and homogenized in 50% trichloroacetic acid in PBS. The lysate was centrifuged at 15,000 rpm min^−1^ twice for 15 min at 4 °C, and the supernatant was used to measure fluorescence (excitation/ emission = 620/680 nm, SpectraMax M2e, Molecular Devices). The relative fluorescence units (R.F.U.) were normalized by the brain weights. To histologically confirm the plasma leakage into the brain parenchyma, 70 kDa dextran-TMR was injected through jugular vein and let circulate for 1h. Brain was removed without perfusion, embedded OCT compound directly and frozen [72]. Sections were cut (9 µm) in a cryostat and fixed with 4% PFA before Immunohistochemistry. After staining with anti Glut-1 antibody, the ipsilateral cortex was observed on an FluoView FV10i confocal microscope (Olympus, Japan) (original magnification, × 60).

### Statistical analysis

All results were expressed as mean ± SEM. Statistical analysis were performed with GraphPad Prism (GraphPad Software, Version 7.0c) by using two-tailed Student’s t-test, one-way analysis of variance (ANOVA) followed by Tukey’s test or two-way ANOVA followed by Bonferoni’s test. *P* values < 0.05 were considered statistically significant.

## RESULTS

### Expression of S1PR1 in the mouse and human brain

We first investigated the expression of S1PR in brain, mouse primary neurons and mouse brain endothelial cells by reverse transcription and quantitative PCR analysis (RT-qPCR), as we have previously described [62, 73]. We found that S1PR1 is robustly expressed, being the most abundant S1PR transcript in the mouse brain *in vivo*, (Fig. 1A, 20.5± 2.4 copies/10^6^ 18S, which is equivalent to approximately 20.5± 2.4 S1PR1 mRNA copies/cell [74]), as well as in the mouse brain endothelial cell line bEnd.3 [75] (Fig. 1B, 87.3±2.9 copies/cell), mouse primary neurons (Fig. 1C, 16.4±1.3 copies/cell) and mouse primary mixed glial cells (Fig. 1D, 8.3±0.15 copies/cell) *in vitro*. Then, we compared the expression of S1PR1 in cortical microvessels fragments [56] to its expression in total brain. Interestingly, we found that S1PR1 transcripts are highly enriched in cortical microvessels when compared to total brain (Fig. 1E, ~6 fold enrichment in microvessels *vs* whole brain).

**Figure 1.**
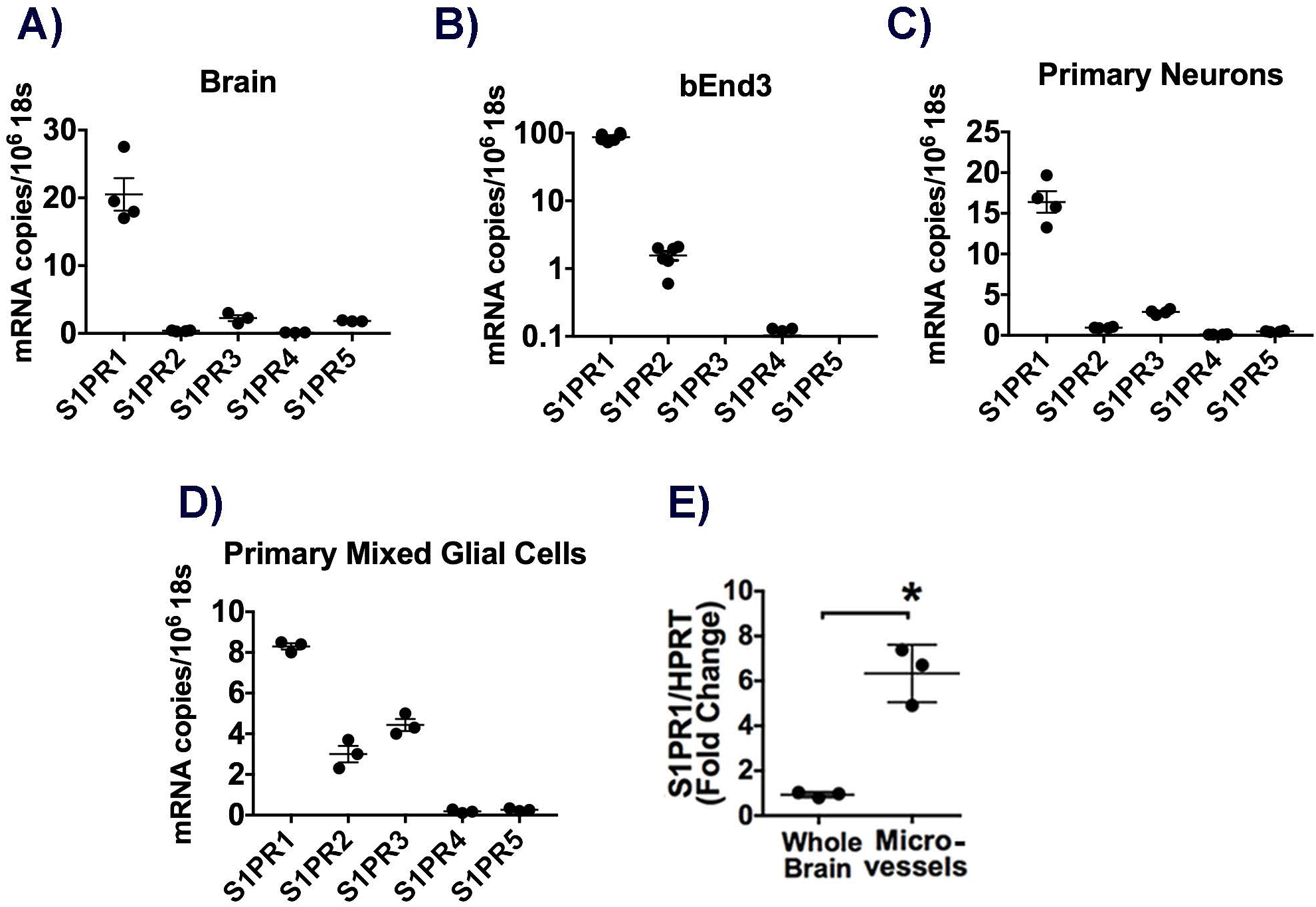
S1PR1 mRNA is the most abundant S1PR transcript in brain, brain endothelial cells and neurons and it is highly enriched in brain microvessels. S1PR mRNA levels in A) normal mouse brain, B) the mouse brain endothelial cell line bEnd.3, C) primary mouse neurons and D) primary mouse mixed glial cells. E) Comparison of S1PR1 mRNA levels in brain microvessels versus whole brain. Quantitative reverse transcription and polymerase chain reaction (RT-qPCR) demonstrates that S1PR1 transcript is predominant over the other S1PR and it is highly enriched in brain microvessels compared to whole brain. B) Log scale. The individual values and the mean ± SEM are shown. N=3-6. *p<0.05.

Using the S1pr1-eGFP knock in mice, in which S1PR1-eGFP fusion protein is expressed under the control of the endogenous S1PR1 promoter [58], we found that S1PR1 is abundantly expressed in the endothelium of cerebral microvessels throughout the brain (Fig. 2A, B, representative pictures from corpus callosum and cortex) as assessed by detection of S1PR1-eGFP and immunofluorescence (IF) analysis for the glucose transporter 1 (Glut-1).

**Figure 2.**
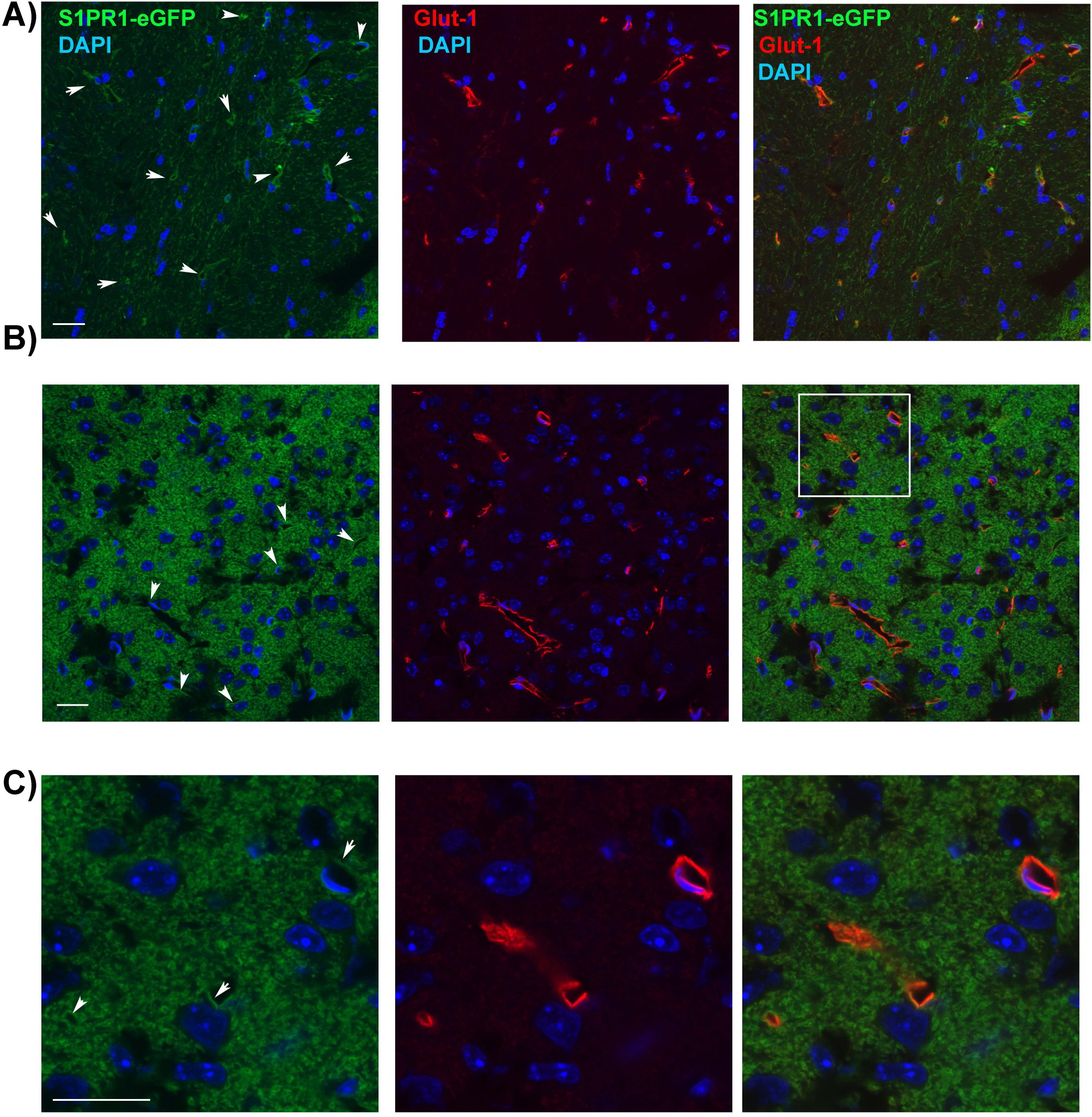
S1PR1 is expressed in the cerebrovascular endothelium of the mouse brain. Confocal analysis of S1PR1-eGFP fluorescence and Glut-1 immunodetection in white matter (A, corpus callosum) and grey matter (B, C, cortex) areas of the mouse brain. S1PR1-eGFP (green channel) is expressed in the cerebrovascular endothelium. Immunofluorescence for Glut-1 (endothelial marker) is shown in the red channel. C) Inset shown in panel (digital zoom). Representative pictures are shown. Scale bar 20μm. N=5-6. Sections were captured using an FluoView FV10i confocal microscope (Olympus, Japan) (original magnification, × 60). Arrows in the green channel point to bright green signal in the microvessels (Glut1 positive).

S1PR1 is also expressed in neurons in various anatomical areas, mainly in the neuropil, as assessed by Nissl and microtubule associated protein (MAP)-2 staining. Representative pictures of cortex and hippocampus are shown (Supplementary Figures 1 and 2, A and B). In white matter areas (e.g. corpus callosum and internal capsule, supplementary figures 1C and 2C, D), S1PR1-eGFP is localized mainly in the neuronal processes, surrounding MAP2 signal. Only some fibrous astrocytes express S1PR1 (Supplementary Figure 3).

A similar pattern of expression was observed in the mouse brain when immunofluorescence analysis was conducted using a S1PR1 antibody previously validated in our laboratory [62] (data not shown).

In order to investigate the expression of S1PR1 in the human brain, we conducted IHC analysis in sections from the frontal lobe. The S1PR1 antibody and IHC protocol used in human samples were previously validated in our laboratory [62]. Representative pictures of the frontal cortex are shown in Figure 3. S1PR1 immuno positivity was observed both in the grey matter (GM) and subcortical white matter (WM) areas (Figure 3A, low magnification). S1PR1 was widely detected in the cerebrovascular endothelium of parenchymal vessels (in the grey matter and white matter, Fig. 3B, D-F) and pial vessels (Fig. 3C). In the grey matter, S1PR1 staining was observed mainly in the neuropil (Fig. 3B, C). In the subcortical white matter, the detection of S1PR1 was more diffuse and weaker compared to the signal observed in the microvessels (Figure 3E and F).

**Figure 3.**
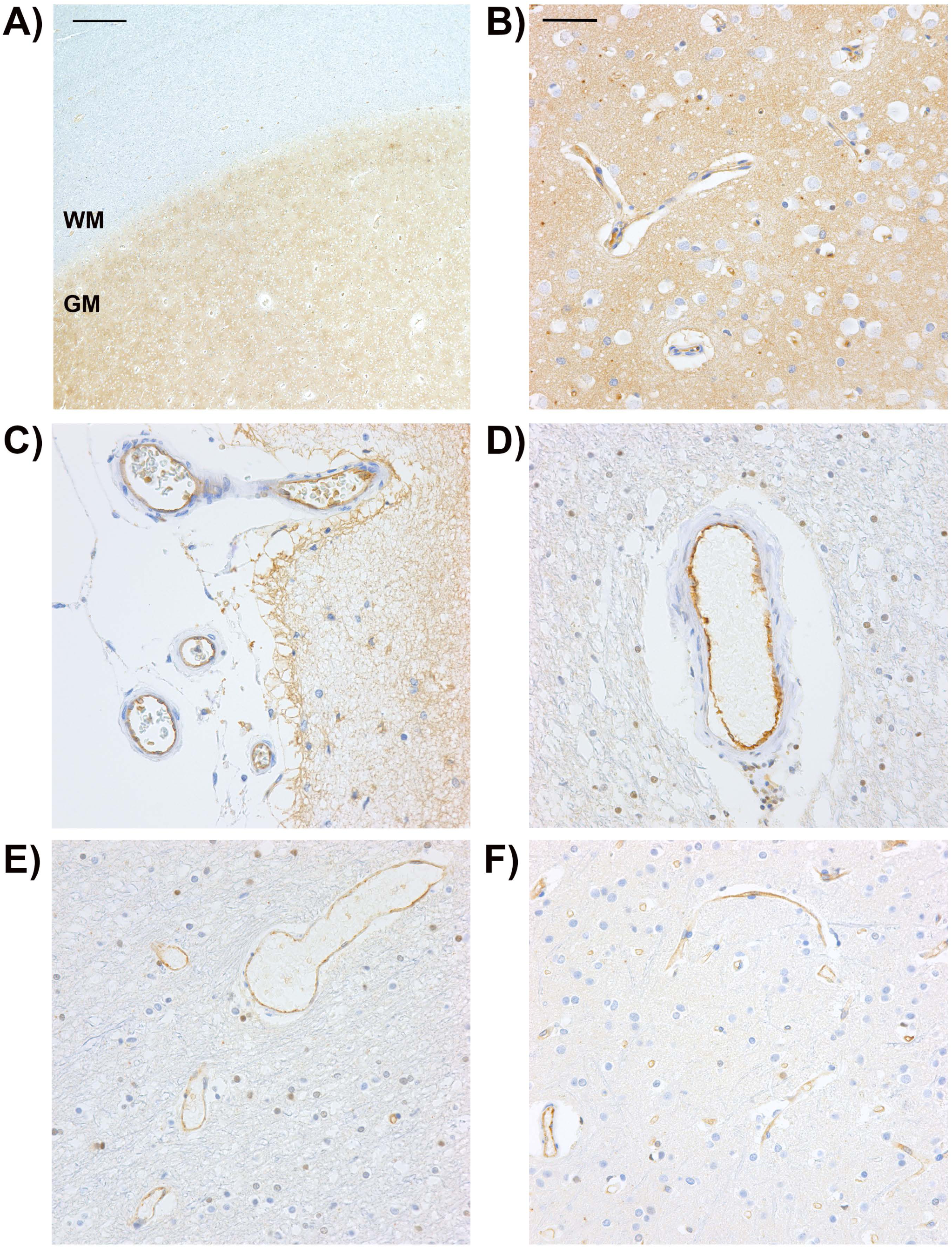
S1PR1 is detected in the human cerebrovascular endothelium. Representative images of S1PR1 immunohistochemistry from 5 human brain autopsy samples. Formalin-fixed paraffin-embedded tissue sections were used for immunohistochemical staining of S1PR1, as described in the Methods section. A) Expression of S1PR1 in frontal cortex grey matter (GM) and subcortical white matter (WM). Scale bar 500 μM. B) Representative picture showing S1PR1 immunopositivity in arterioles and capillaries as well as the neuropil of the cortical grey matter. C) Detection of S1PR1 in the cerebrovascular endothelium of pial vessels. D-F) Representative pictures of S1PR1 immunodetection in the subcortical white matter. Notice S1PR1 positivity in parenchymal arterioles (D, F), venules (E) and capillaries (E, F). B-F) Scale bar 50 μM. Pictures were taken with a 60x objective (Olympus, Japan).

Altogether, our data indicate that, S1PR1 is the most abundant S1PR transcript in the brain, and in brain endothelial cells, neurons and mixed glial cells *in vitro*. S1PR1 protein is also widely detected in the mouse and human brain, in the cerebrovascular endothelium and the grey matter neuropil. S1PR1 mRNA is highly enriched in mouse cerebral microvessels when compared to total brain.

### Endothelial S1PR1 is an endogenous protective pathway induced in cerebral microvessels after SAH

In order to investigate the role of endothelial S1PR1 signaling in the pathophysiology of aneurysmal SAH, we used the endovascular rupture model of SAH[61], which is a well established and reproducible model that recapitulates key features of the pathophysiology of the acute phase of SAH. 24, 48 and 72 hours after SAH, cerebral microvessels were isolated from the mouse cortex of wild type mice to determine sphingosine 1-phosphate receptor 1 (S1PR1) mRNA and protein levels compared to sham mice. We found that S1PR1 mRNA levels in cortical microvessels were significantly increased at 24h after SAH (3.14±0.55 fold induction) compared to sham (Fig. 4A). Gfap mRNA levels in microvessels were significantly higher 24 h after SAH compared to sham (Fig. 4B, 5.14± 0.55 fold induction). The levels of endothelial (Tjp-1, Fig. 4C), pericyte (Anpep, CD13, Fig. 4D) or astrocyte end foot (Aqp4, Fig. 4E) markers did not significantly change suggesting similar cell composition among the microvessel preparations from these groups of mice. S1PR1 protein was widely detected by western blot assay in cerebral microvessels in sham animals (Supplementary Figure 4). A modest increase in S1PR1 protein levels at 24 hours after SAH was observed (1.68 ± 0.19-fold). These data indicate that, in the acute phase of SAH, neurovascular inflammation occurs. In addition, S1PR1 mRNA and protein levels in cerebral microvessels were significantly increased upon SAH.

**Figure 4.**
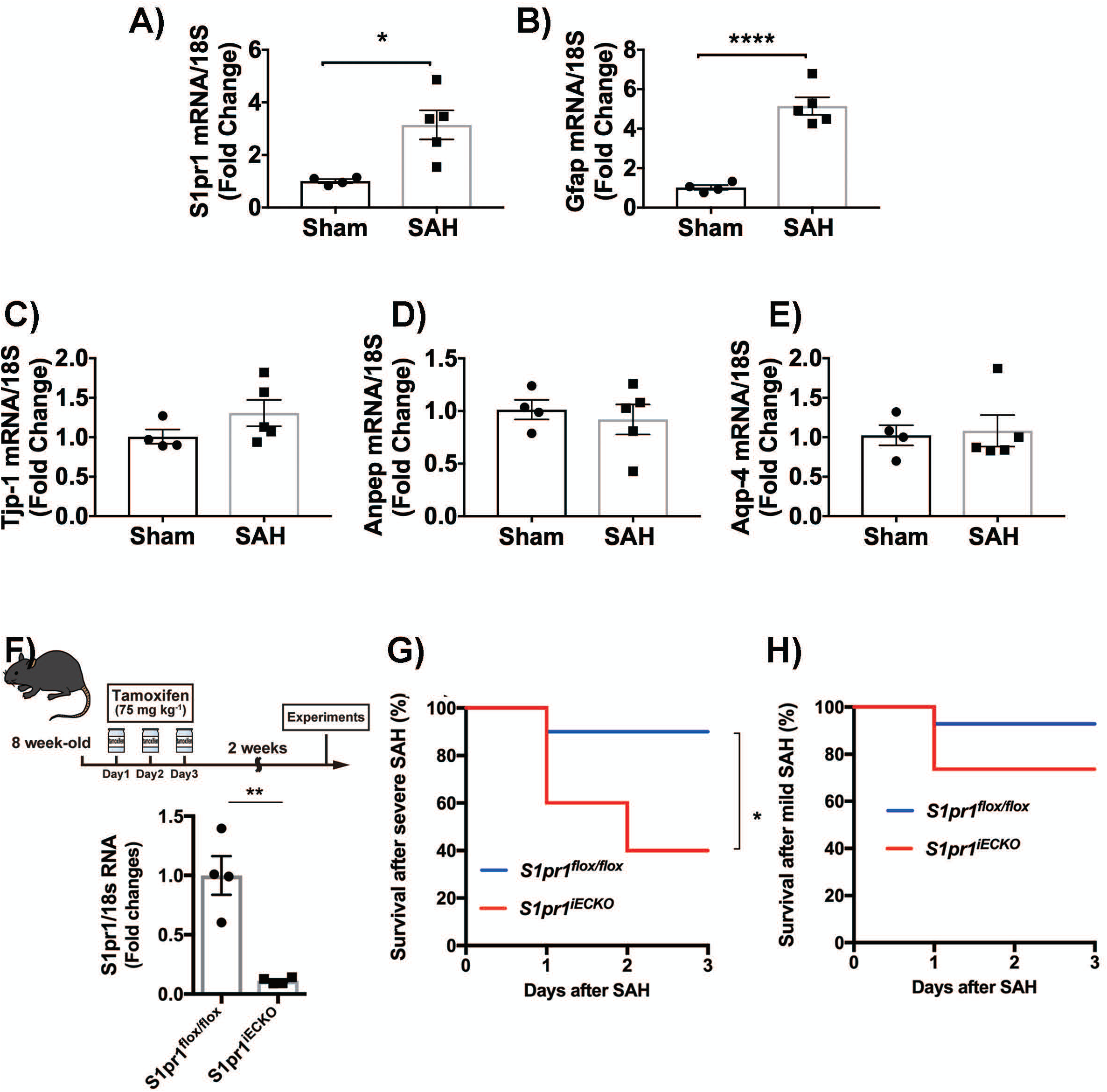
Endothelial S1PR1 is an endogenous protective pathway induced in cerebral microvessels after SAH. A-E) S1pr1, Gfap, Tjp-1 (ZO-1), Anpep (CD13) and Aqp4 mRNA levels in cerebral microvessels were quantified by RT-qPCR in sham animals 24 h after SAH. S1pr1 and Gfap are induced in cerebral microvessels after SAH. The mRNA levels of the endothelial (ZO-1, Tjp-1), pericyte (Anpep, CD13) or astrocyte (Aqp4) markers did not significantly change suggesting similar cell composition among the microvessel preparations from these groups of mice. F) Efficiency of deletion of S1PR1 in the cerebrovascular endothelium. *S1pr1^flox/flox^* and *S1pr1^iECKO^* mice were treated with tamoxifen (75 mg kg^−1^) for the consecutive 3 days at the age of 8 weeks. *S1pr1* mRNA levels were analyzed by qPCR in brain endothelial cells isolated from *S1pr1^flox/flox^* and *S1pr1^iECKO^* mice, 2 weeks after tamoxifen treatment (*n* = 4). Efficiency of deletion in the cerebrovascular endothelium in *S1pr1^iECKO^* mice is shown relative to *S1pr1^flox/flox^* mice. Individual values and mean ± SEM are shown. \**P* < 0.05, \*\**P* < 0.01, \*\*\**P* < 0.001, Student’s t-test. G) Survival curves in *S1pr1^flox/flox^* and *S1pr1^iECKO^* mice after severe SAH surgery (*n* = 10). H) Survival curves in *S1pr1^flox/flox^* and *S1pr1^iECKO^* mice after mild SAH surgery (*n* = 13 or 19). G and H) Log-rank test, \**P* < 0.05. Red line, *S1pr1^iECKO^*; blue line, *S1pr1^flox/flox^*.

Given the abundant expression of S1PR1 in the cerebrovascular endothelium in the mouse and human brain, and the enrichment in S1PR1 mRNA in mouse cortical microvessels compared to total brain, we aimed to investigate the role of endothelial S1PR1 signaling in early brain injury (EBI) after SAH. We generated endothelial-specific S1PR1 null mice (S1PR1 ^iECKO^) by treating adult *S1pr1^flox/flox^xCdh5–Cre^ERT2^* mice (~2 month old) with 75 mg kg^−1^ tamoxifen for 3 consecutive days (Fig.4F) as we have recently described[56]. *S1pr1^flox/flox^* littermate mice were also treated with tamoxifen and used as wild type control. We assessed the efficiency of deletion in S1PR1^iECKO^ by isolating brain endothelial cells from mice 2 weeks after tamoxifen treatment. RT-qPCR analysis demonstrated ~90% reduction of *S1pr1* expression in *S1pr1^iECKO^* mice compared to *S1pr1^flox/flox^* littermate mice treated with tamoxifen in the same manner (Fig. 4F). We previously reported that, in resting conditions, postnatal endothelial deletion of S1PR1 does not have an impact on central nervous system vascular development, maturation, or pattern formation [56].

*S1pr1^iECKO^* and *S1pr1^flox/flox^* littermates were subjected to SAH surgery using a modified 5.0 suture (0.3 mm × 1.7 mm) to induce severe SAH as described in the method section [61]. The mortality rates were 10% in *S1pr1^flox/flox^* mice at day 1, 2 and 3 after SAH. Interestingly, the mortality rates in endothelial-specific S1PR1 null mice (*S1pr1^iECKO^*, 40% at day 1, and 60% at day 2 and 3, Fig. 4G) were significantly higher compared to wild type (*S1pr1^flox/flox^*,.

Given the high mortality rate in *S1pr1^iECKO^* subjected to the severe SAH model (Fig. 4G), we used a milder SAH model for subsequent studies, by changing the shape of the tip of the nylon suture to perforate the cerebral artery (0.3 mm × 0.3 mm, as described in methods section). When *S1pr1^iECKO^* mice were challenged with mild SAH, they exhibited a trend towards a higher mortality rate (26.3% in day 1, 2 and 3) compared to *S1pr1^flox/flox^*, which exhibited a mortality rate of 7.7% at day 1, 2 and 3 (Fig. 4H), although it was not statistically significant. Altogether these data indicate that endothelial-specific deletion of S1PR1 in mice increases mortality in the acute phase of SAH.

### Genetic deletion of *S1pr1* in the endothelium has no effect on blood volume in the subarachnoid space, hemostasis or CBF changes after SAH

The volume of blood in the subarachnoid space, which depends on the amount of bleeding and the clearance, directly correlates with worse outcomes after SAH [76] [63]. Thus, we quantified the amount of subarachnoid blood upon SAH in *S1pr1^flox/flox^* and *S1pr1^iECKO^* mice by image analysis using a previously described SAH grading system [63]. No significant differences were observed in the amount of subarachnoid blood between *S1pr1^flox/flox^* and *S1pr1^iECKO^* mice 24h after SAH (Fig. 5A, representative pictures). SAH grading was 11.50 ± 0.62 in *S1pr1^flox/flox^* mice and 11.67 ± 0.42 in *S1pr1^iECKO^* (Fig. 5B). In addition, we determined the role of endothelial-specific S1PR1 in hemostasis using the tail bleeding assay [64]. We did not find any significant differences in the bleeding times or blood volumes between *S1pr1^flox/flox^* and *S1pr1^iECKO^* mice in the tail bleeding assay. Bleeding times were 68.88 ± 4.98 seconds in *S1pr1^flox/flox^* and 63.8 ± 4.14 seconds in *S1pr1^iECKO^* (Fig. 5C). Hemoglobin contents, assessed by measuring the absorbance at 550 nm, were 0.73 ± 0.07 in *S1pr1^iECKO^* mice compared to 0.70 ± 0.12 in *S1pr1^flox/flox^* mice (Fig. 5D). Altogether, these data indicate that hemostasis is not impaired in *S1pr1^iECKO^* mice compared to wild type mice and that the amount of blood in the subarachnoid space 24 h upon SAH is similar in wild type and *S1pr1^iECKO^ mice*.

**Figure 5.**
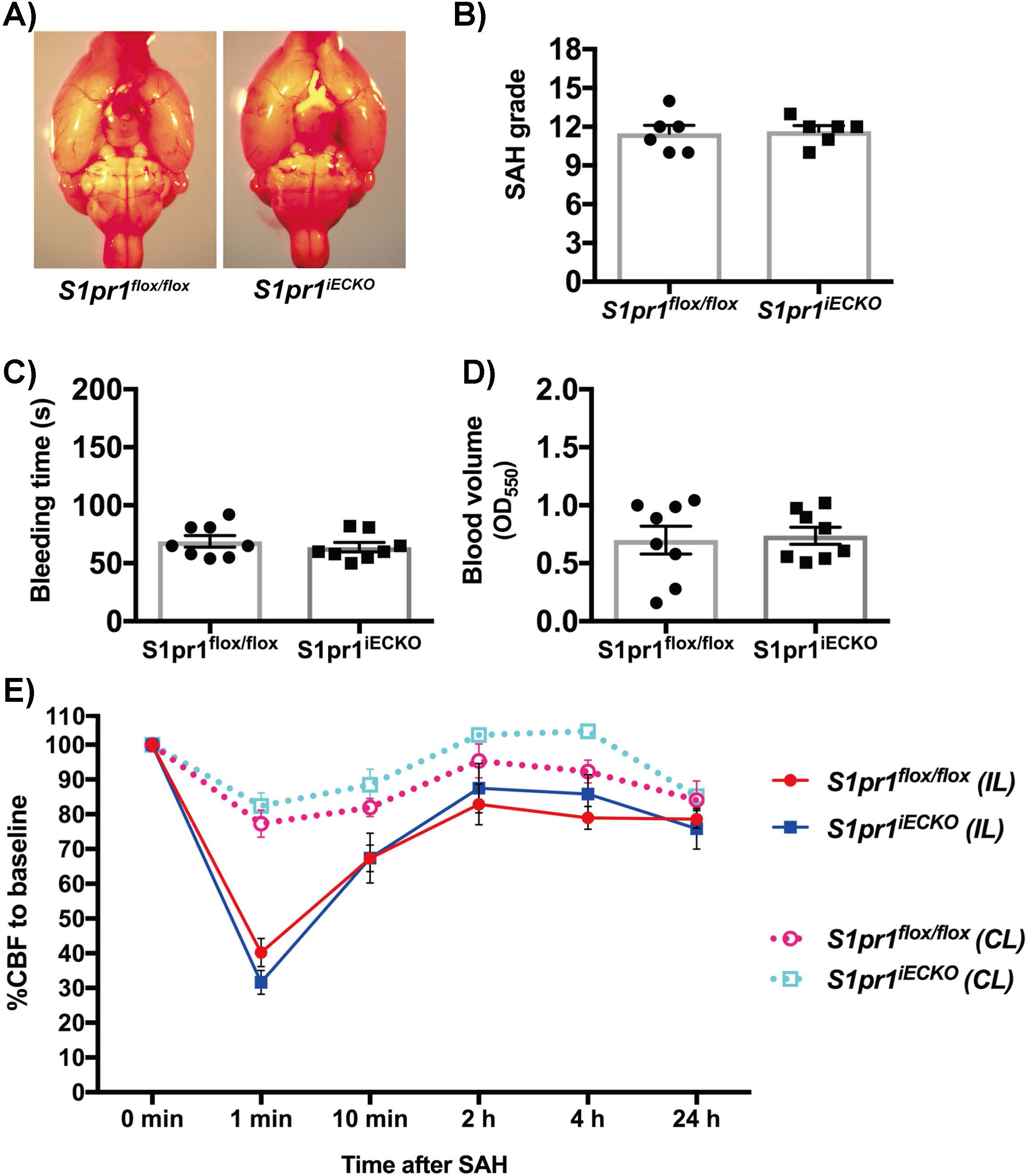
Genetic deletion of *S1pr1* in the endothelium has no effect on subarachnoid blood volume, hemostasis or CBF changes after SAH. A) Representative images of the ventral side of the brain from *S1pr1^flox/flox^* and *S1pr1^iECKO^* mice at 24 hours after SAH. B) Subarachnoid blood volume (SAH grade) was calculated by quantifying the blood in the subarachnoid space by image analysis as described in methods section. (*n* = 6). (C-D) Quantification of hemostasis in the tail bleeding assay. C) bleeding time and D) blood volume (*n* = 8). (B-D) Student’s t-test. Individual values and mean±SEM are shown. (E) CBF in the middle cerebral artery territory was measured by Laser-speckle contrast imager in *S1pr1^flox/flox^* and *S1pr1^iECKO^* mice before (0 min.), 1 min, 10 min, 2 h, 4 h, 24 h after SAH induction. The relative CBF values (%) versus before SAH are shown. Data are mean ± SEM. (*n* = 6). Two-way ANOVA followed by Bonferoni’s test showed no statistically significant differences between wild type and *S1PR1^iECKO^*. Red solid line, *S1PR1^flox/flox^* ipsilateral side; blue solid, *S1PR1^iECKO^* ipsilateral; light dotted line, *S1PR1^flox/flox^* contralateral; light blue dotted line, *S1PR1^iECKO^* contralateral. IL, ipsilateral side. CL, contralateral side.

We also determined cerebral blood flow (CBF) changes during SAH in both groups of mice by Laser-Speckle flowmetry (Figure 5E). CBF in the middle cerebral artery (MCA) territory rapidly dropped ~1 minute after SAH in a similar way in both *S1pr1^flox/flox^* and *S1pr1^iECKO^* mice. In the ipsilateral hemisphere, CBF dropped to 40.26 ± 4.09% of basal in *S1pr1^flox/flox^ and* to 31.65 ± 3.46% of basal in *S1pr1^iECKO^*; in the contralateral hemisphere, CBF dropped to 77.32 ± 3.90% of basal in *S1pr1^flox/flox^* and to 82.42 ± 3.72% of basal in *S1pr1^iECKO^*. Afterwards, as shown in Figure 5E, CBF progressively recovered similarly in S*1pr1^flox/flox^* and *S1pr1^iECKO^* mice. 2 h after SAH, in the ipsilateral hemisphere, CBF recovered to 82.83 ± 5.88% of basal in *S1pr1^flox/flox^ and* to 87.46 ± 7.15% of basal in *S1pr1^iECKO^*; in the contralateral side CBF recovered to 95.34 ± 4.86% of basal in *S1pr1^flox/flox^* and to 102.81 ± 1.97% of basal in *S1pr1^iECKO^*. These data indicate that genetic deletion of *S1pr1* in the endothelium did not have a significant impact in CBF changes upon SAH.

Finally, no significant differences were observed in arterial O_2_ saturation, heart rate, pulse distention (a surrogate of pulse pressure) and respiratory rate between *S1pr1^iECKO^* and *S1pr1^flox/flox^* mice, before or after SAH (Table 1).

Altogether, these data indicate that endothelial-specific deletion of S1PR1 in adult mice did not have a significant impact on bleeding or clearance of subarachnoid blood, cerebral blood flow changes or systemic physiological parameters such as heart rate, respiratory rate, O_2_ saturation or pulse distension.

### Endothelial deletion of *S1pr1* exacerbates brain edema and cell death after SAH leading to worsened neurological outcomes

In order to determine the impact of the lack of endothelial S1PR1signaling on brain injury after SAH, we assessed brain edema at 72 hours after induction of mild SAH by quantifying total brain water content. *S1pr1^flox/flox^* mice exhibited a statistically significant increase in brain water content in the ipsilateral hemisphere (82.83 ± 0.53%) compared to sham (81.34 ± 0.20%) (Fig. 6A). Interestingly, in *S1pr1^iECKO^* mice, total brain edema in the ipsilateral hemisphere was significantly exacerbated (84.39 ± 0.91%) compared to *S1pr1^flox/flox^*(Fig. 6A). There were no changes in brain water content in contralateral hemisphere (SAH versus sham) in either *S1pr1^flox/flox^ or S1pr1^iECKO^* mice.

**Figure 6.**
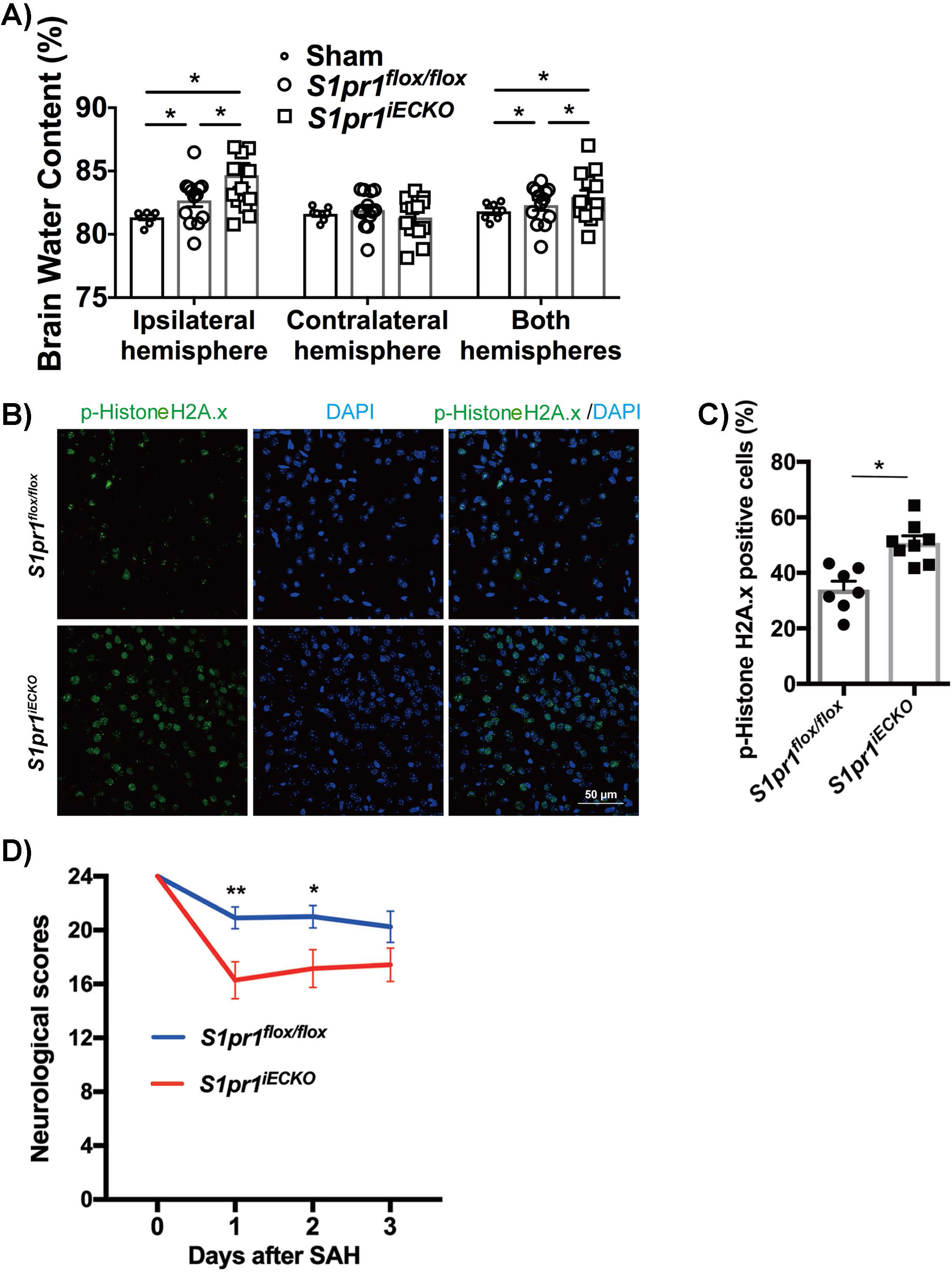
Exacerbated cerebral edema and neuronal injury after SAH in endothelial specific S1pr1 null mice. A) 72 hours after SAH, brain edema was evaluated by quantification of brain water content in *S1pr1^flox/flox^* and *S1pr1^iECKO^* mice (*n* = 7 - 14). Brain water content in the ipsilateral and contralateral hemispheres was calculated as ([wet weight-dry weight]/wet weight) × 100. * p<0.05, one-way ANOVA followed by Tukey’s test. B) Immunofluorescence confocal analysis of phospho-histone H2A.X, a marker for DNA damage (green channel). Nuclear staining (DAPI) is shown in the blue channel. Representative images of ipsilateral hemisphere (cortex) from *S1pr1^flox/flox^* and *S1pr1^iECKO^* mice after SAH stained are shown. Scale bar, 50 µm. C) Quantification of phospho-histone H2A.X positive cells (%) (n = 7-8 mice). \**P* < 0.05, student’s t-test. The individual values and the mean ± SEM are shown. Each data point represents a mouse and it is the average of 3 different fields. D) Neurological (sensory and motor) deficits after mild SAH surgery in *S1pr1^flox/flox^* and *S1pr1^iECKO^* mice (n = 12 or 14) were assessed as described in methods. From 4 to 24 points: 24 points (best), 4 points (worst). Data are mean ± SEM. \**P* < 0.05, \*\**P* < 0.01, One-way ANOVA followed by Tukey’s test. Red line, *S1pr1^iECKO^*; Blue line, *S1pr1^flox/flox^*.

Next, we analyzed cell death at 24 hours after SAH using phospho-histone H2A.X (Ser 139) immunostaining, a marker for apoptosis and DNA damage [69, 70]. We found that *S1pr1^iECKO^* mice showed significantly higher number of phospho-histone H2A.X positive cells compared to *S1pr1^flox/flox^* mice (50.85 ± 2.56% in *S1pr1^iECKO^* versus 34.00 ± 2.98% in *S1pr1^flox/flox^*) (Fig. 6B and C). No apoptotic cells were detected in sham animals.

Finally, we aimed to determine the impact of genetic deletion of S1PR1 in the endothelium on neurological outcomes after SAH. Neurological outcomes were determined by assessing motor and sensory function using a total scale of 4 to 24 (being 24 the best neurological outcome) as previously described[65, 66]. Neurological outcomes at 24h and 48h after mild SAH surgery were worsened in *S1pr1^iECKO^* mice (16.29 ± 1.37 and 17.14 ± 1.40, respectively) compared to *S1pr1^flox/flox^* mice (20.91 ± 0.81 and 21.00 ± 0.83, respectively) (Fig. 6D).

These data indicate that genetic deletion of *S1pr1* specifically in the endothelium significantly exacerbates total brain edema and cell death resulting in poorer neurological outcomes.

### Endothelial specific deletion of *S1pr1* in adult mice exacerbates blood brain dysfunction after SAH

Given the critical role of the cerebrovascular endothelium in BBB maintenance, we determined BBB dysfunction 24 hours after SAH or sham surgery. Albumin leakage was quantified by Evans Blue Dye (EBD) extravasation assay, in *S1pr1^flox/flox^* and *S1pr1^iECKO^* mice. In addition, leakage of macromolecules into the brain parenchyma was histologically confirmed by detection of intravenously injected 70kDa tetramethylrhodamine (TMR)-dextran and immunofluorescence analysis of the endothelial marker Glut-1. We found that 24 h after SAH, albumin leakage in the ipsilateral hemisphere was significantly increased in *S1pr1^flox/flox^* mice compared to sham animals (Fig 7A, 1.53 ± 0.06 fold in *S1pr1^flox/flox^* SAH versus *S1pr1^flox/flox^* sham) and significantly higher in *S1pr1^iECKO^* mice with regards to *S1pr1^flox/flox^*(1.83 ± 0.08 in *S1pr1^iECKO^*, Fig. 7A). In addition, upon sham surgeries, no differences in albumin leakage were observed between *S1pr1^flox/flox^* and *S1pr1^iECKO^*, consistent with our recent report [56]. Histological analysis of 70kDa tetramethylrhodamine (TMR)-dextran localization, confirmed increased leakage into the brain parenchyma and outside of cerebral capillaries upon SAH (Fig. 7B). Altogether, these data indicate that BBB dysfunction upon SAH is exacerbated in animals lacking S1PR1 specifically in the endothelium.

**Figure 7.**
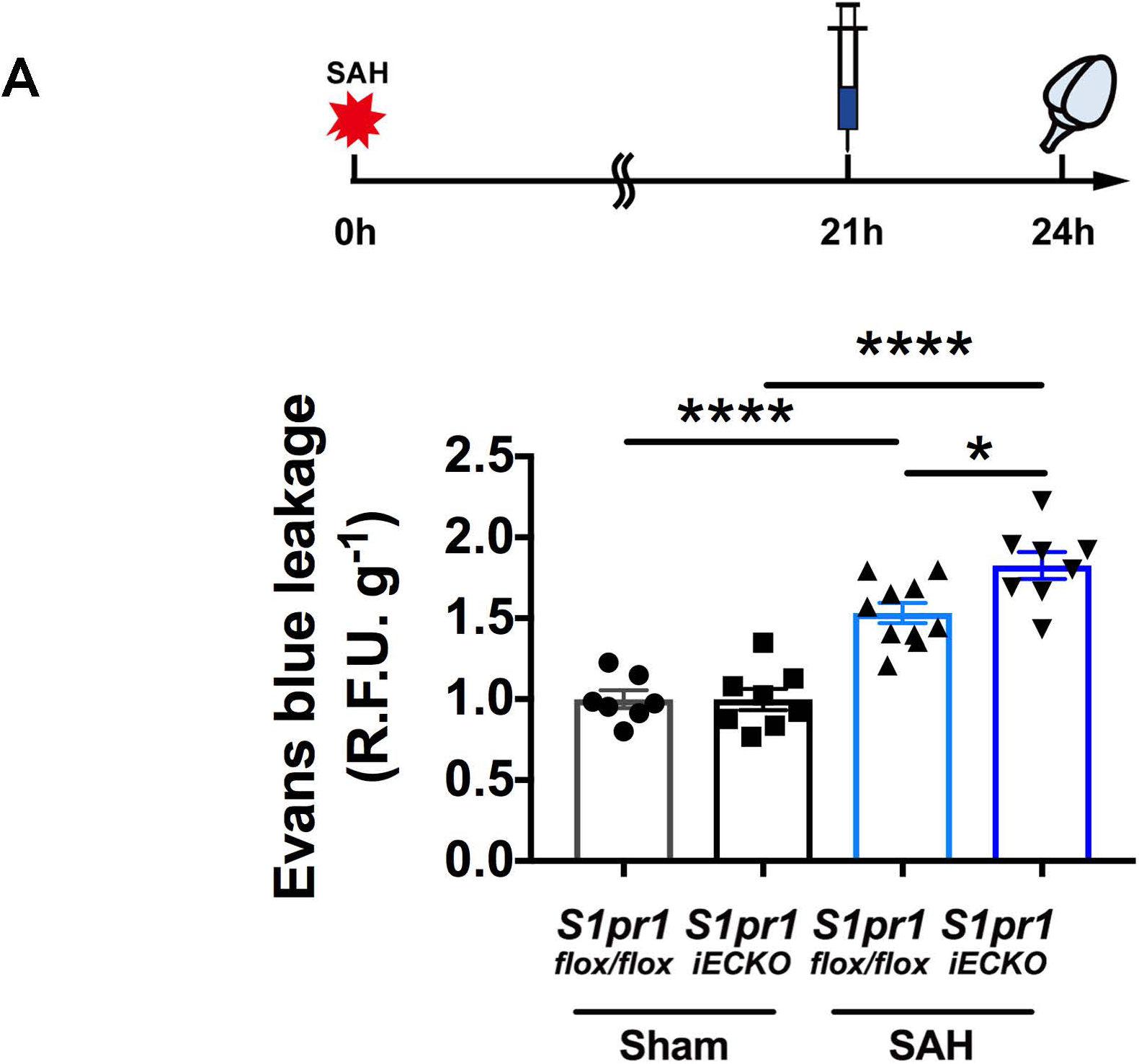
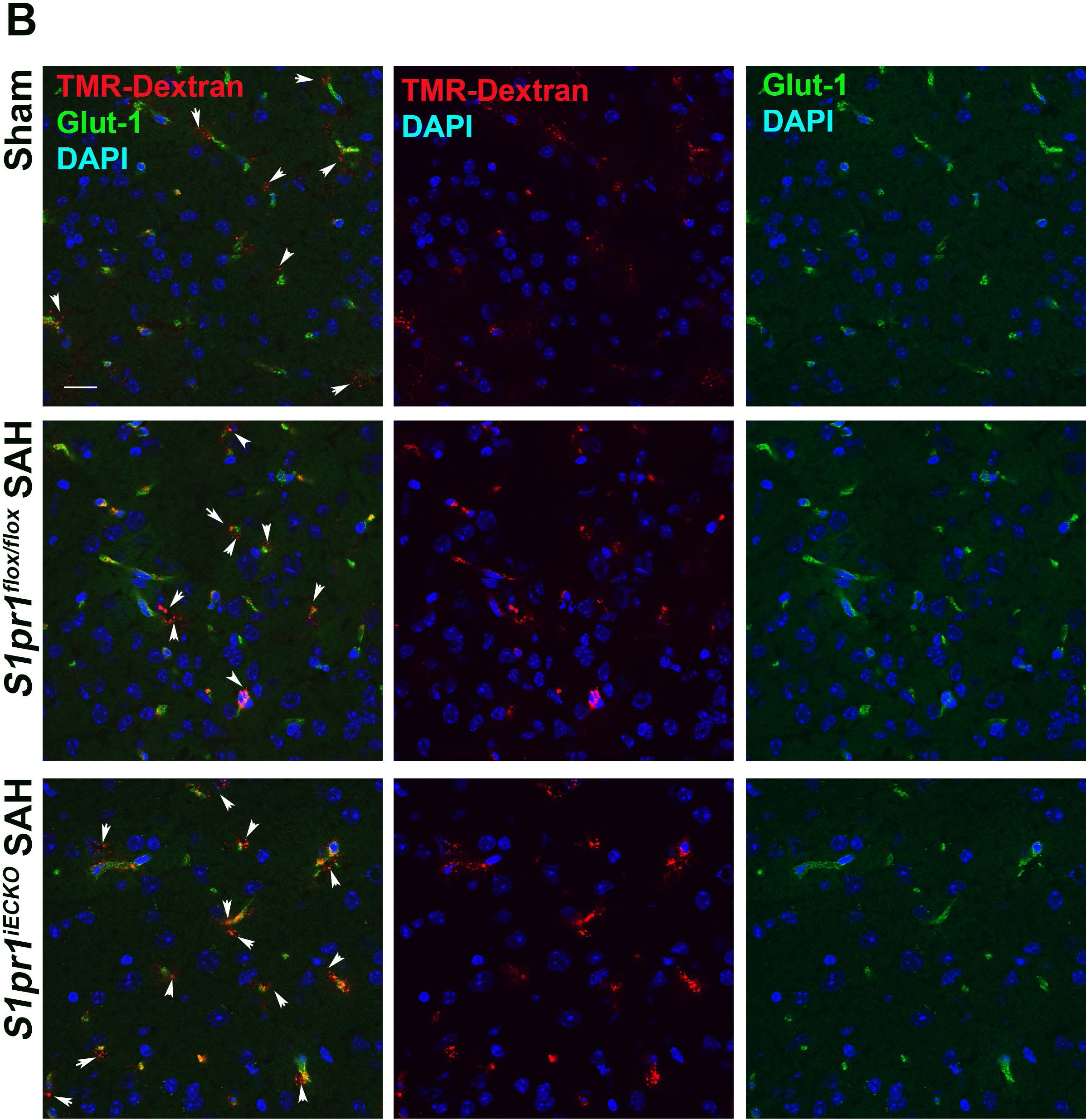
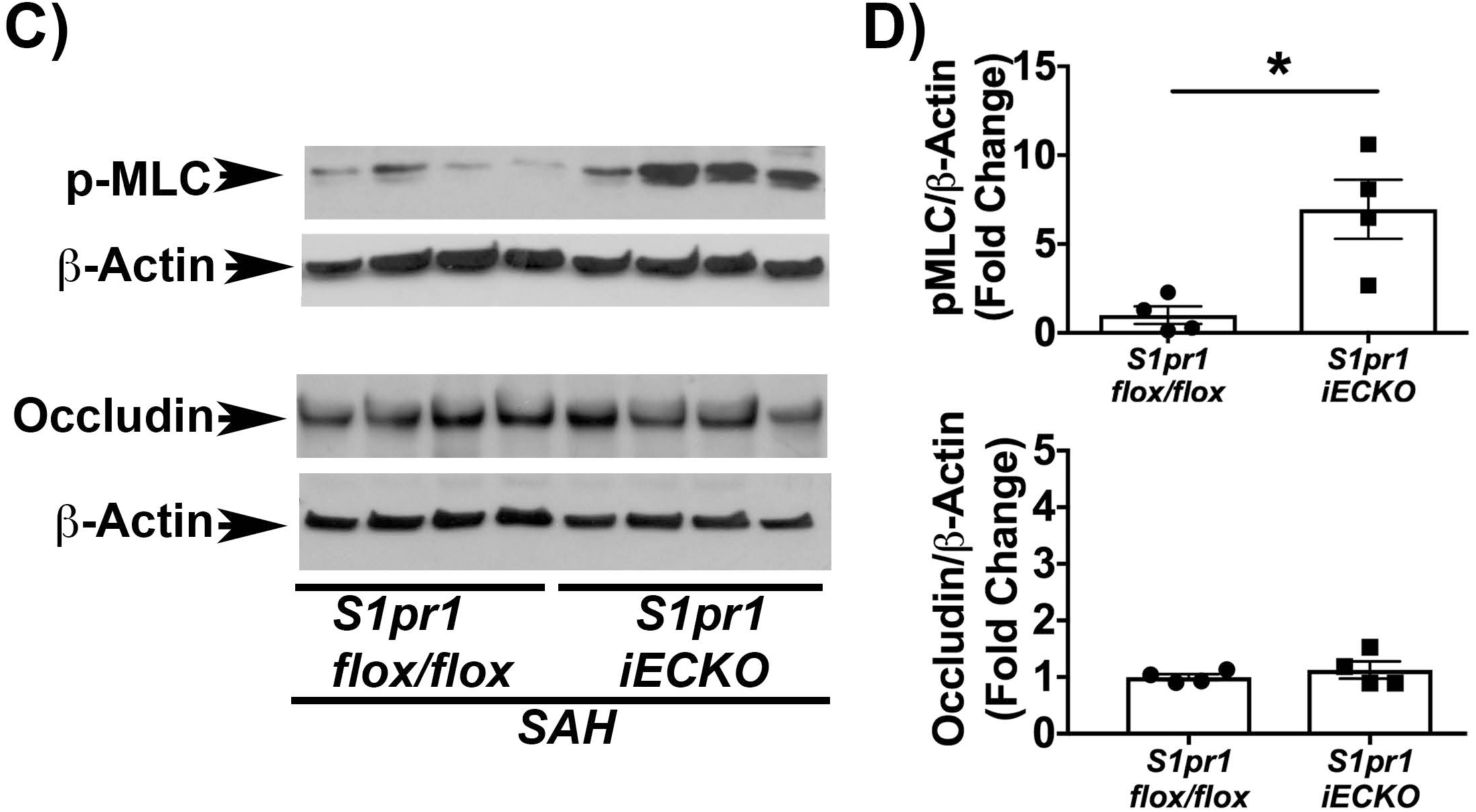
Endothelial specific S1pr1 null mice exhibit exacerbated BBB leakage after SAH compared to wild type littermates. A) Albumin BBB leakage, assessed by Evans Blue Dye extravasation, 24 hours after sham or SAH surgery in *S1pr1^flox/flox^* and *S1pr1^iECKO^* mice (*n* = 6 - 10). Evans blue dye was circulating for 3 hours. The individual values and mean ± SEM are shown. \**P* < 0.05, \*\**P* < 0.01, \*\*\**P* < 0.001, \*\*\*\**P* < 0.0001 (One-way ANOVA followed by Tukey’s test). *y* axis shows relative fluorescence units (R.F.U.) per gram of tissue. B) Histological determination of 70kDa of TMR-dextran leakage (red channel) into the brain parenchyma. Immunofluorescence for Glut-1 (endothelial marker) is shown in the green channel. Nuclear stain (DAPI) is shown in the blue channel. White arrows point to extravascular-parenchymal TMR-dextran. Representative pictures are shown. Scale bar 20μm. N=5-6. Sections were captured using an FluoView FV10i confocal microscope and a 60x objective (Olympus, Japan). C-D) Increased levels of phosphorylated myosin light chain (p-MLC) in isolated cortical microvessels in *S1pr1^iECKO^* mice compared to *S1pr1^flox/flox^*. Cortical microvessels were isolated 24 h after SAH in *S1pr1^iECKO^* and *S1pr1^flox/flox^* mice. C) Western blot analysis for phospho-MLC (p-MLC), occludin and β-actin. D) Western blot quantification was conducted using Image J. Individual values and mean ± SEM are shown. \**P* < 0.05 (t test).

In order to shed light on the mechanisms whereby endothelial deletion of S1PR1 leads to exacerbated BBB dysfunction, we isolated cerebral microvessels after SAH from *S1pr1^flox/flox^ S1pr1^iECKO^* mice and determined the activation of signaling pathways implicated in BBB dysfunction by western blot analysis. The Rho-ROCK pathway has been shown to play a critical role in endothelial barrier dysfunction, endothelial inflammation [77] and also in blood brain barrier dysfunction in central nervous system pathologies [78–80]. We found that exacerbated BBB dysfunction in endothelial-specific S1PR1 null mice correlated with increased phosphorylation of myosin light chain (MLC) in isolated cortical microvessels (Fig. 7C and D, 6.96±1.6 fold in *S1pr1^iECKO^ versus S1pr1^flox/flox^*), a downstream effector of the Rho-ROCK pathway critical for the induction blood brain barrier dysfunction [78]. The levels of endothelial markers (e.g. occludin, Fig. 7C and D) were similar, indicating consistent cell composition between microvessel preparations [59]. Altogether, these data indicate that BBB dysfunction after SAH (albumin and 70 KDa dextran leakage) is heightened in mice lacking S1PR1 specifically in the endothelium and correlates with increased levels of phosphorylated MLC in cerebral microvessels.

## DISCUSSION

In this study, we aimed to determine the expression of S1PR1 in the cerebrovascular endothelium in mice and humans and the role of endothelial S1PR1 signaling in stroke outcomes in mice. Using tissue specific S1PR1 null mice in which genetic deletion of S1PR1 in the endothelium was induced in the adult and a mouse model of aneurysmal SAH, the most devastating type of stroke, we unveil the critical role of this endothelial pathway in SAH outcomes. We found that S1PR1 transcripts are significantly enriched (~6 fold) in mouse cortical microvessels compared to total brain and that S1PR1 mRNA is induced in cerebral microvessels after SAH. Endothelial specific deletion of S1PR1 resulted in aggravated brain injury and significantly worsened outcomes, indicating that S1PR1 is an endogenous protective mechanism of the endothelium to mitigate exacerbation of neurovascular injury in stroke. *Our study unveiled a previously unappreciated role of the endothelial S1P pathway in the pathophysiology of stroke and implies that activation of this endogenous protective endothelial pathway could have important therapeutic applications in CNS pathologies*.

Cerebral microvascular dysfunction has been implicated in the pathophysiology of numerous acute [5–13, 81] [82–85], and chronic neurological conditions [86–89]. Aneurysmal SAH, the most devastating type of stroke, occurs when an intracranial aneurysm ruptures. Compared to other types of stroke, SAH occurs earlier in life (40-60 years)[90] and leads to higher mortality (50%) [91]. In addition, SAH survivors experience a high degree of disability and cognitive impairment (memory, language and executive function) [92] with personal and societal consequences. SAH causes transient cerebral ischemia and accumulation of blood in the subarachnoid space, leading to brain injury, BBB dysfunction, cytotoxic and vasogenic cerebral edema in the acute phase [12, 93, 94]. Although numerous studies have reported *the direct correlation between endothelial barrier dysfunction and worsened SAH outcomes in humans* [95, 96] and mice [94, 97–102], *the impact of specific endothelial signaling pathways on brain injury and SAH outcomes has remained uncertain*. Given the emerging pathophysiological relevance in humans of the S1PR1 pathway, in this study, we aimed to investigate the role of the endothelial-specific S1PR1 signaling in brain injury in the acute phase of SAH. We used a well-established mouse model of aneurysmal SAH, the endovascular rupture model [61]. This experimental model faithfully recapitulates key features of the acute phase of SAH: upon rupture of the MCA, blood pours into the subarachnoid space, increasing intracranial pressure which gives rise to a brief period of transient (~3-5 minutes) cerebral ischemia [61] (Fig. 5E, ~70%reduction of CBF in ipsilateral hemisphere). CBF slowly recovers over the time, but the brain remains hypoperfused in the acute phase of SAH (Fig. 5E, ~30% reduction vs basal). BBB breakdown, cerebral edema and neuronal injury ensue [97, 100]. Our *study demonstrates the critical role of the endothelium in SAH outcomes and highlights the therapeutic potential of the endothelial S1PR1 pathway to prevent exacerbation of brain injury in this devastating condition*.

There is mounting evidence in humans of the pathophysiological relevance of the S1P-S1PR1 pathway in endothelial [31, 32] and lymphocyte function [38, 39] [40]. Fingolimod [38, 39] and Siponimod [40], 2 immunosupressor drugs FDA approved for the treatment of multiple sclerosis, are pharmacological modulators of S1PR1. Remarkably, they are currently being tested in stroke clinical trials. In experimental stroke, FTY720 and S1PR1 specific agonists, as other immunosuppressor drugs, have been shown to be protective[45, 46] [47, 48]. Although the mechanisms are not completely clear, FTY720 protection has been attributed to its immunosuppressive effects [48] and its ability to desensitize and inhibit S1PR1 signaling. *Due to the immuno-suppressor effects of S1PR1 pharmacological modulators, these previous studies could not determine the role of endothelial specific S1PR1 signaling in BBB function and stroke outcomes*. We sought to address this pivotal question by inducing the deletion of S1PR1 in adult mice, specifically in the endothelium [57] and directly testing the role of endothelial S1PR1 signaling in BBB function and its impact on early brain injury upon SAH. Our data demonstrate that *endothelial S1P signaling via S1PR1 is an endogenous protective signaling pathway that mitigates neurovascular ischemic-hypoxic injury*. These data are consistent with previous studies from our laboratory which indicated that FTY720 exerts agonistic activity for S1PR1 in the endothelium and promotes endothelial barrier function [17, 27]. Overall, our findings with the endothelial-specific S1PR1 mice highlight the therapeutic potential of this endothelial pathway in stroke. *Future strategies to activate this pathway specifically in the endothelium without compromising the immune response hold promise as novel vasoprotective therapeutic agents in stroke*.

Mechanistically, we found that endothelial deletion of S1PR1 exacerbates BBB dysfunction (albumin and 70kDa dextran leakage) upon SAH. BBB dysfunction heightens neurovascular injury by various mechanisms such as by allowing the entrance of neurotoxic plasma components into the brain parenchyma (e.g. albumin [103], fibrinogen [104]) and by compromising the ability of cerebral capillaries to deliver oxygen and nutrients to the neurons [7–9, 105]. In homeostatic conditions, the cerebrovascular endothelium, in cooperation with pericytes and astrocytes maintain a physical, metabolic and transport barrier to restrict the passage of molecules into the brain parenchyma [4, 106, 107]. Crosstalk between the cellular components of the blood brain barrier are critical to maintain the unique phenotype of the cerebrovascular endothelium, characterized by the presence of TJ as well as low expression of molecules involved in vesicular trafficking (e.g. plasmalemma vesicle-associated protein) [108] [3] [109] and leukocyte adhesion (e.g. ICAM-1), which prevents trafficking of plasma molecules and blood cells via paracellular and transcellular routes. Upon ischemic or hypoxic injury, BBB dysfunction ensues, primarily due to two cellular mechanisms, transcellular and paracellular permeability. Gelatinase activity in cerebral microvessels [94, 100, 101, 110–113] is induced leading to basal lamina degradation [7, 114], weakening of pericyte-endothelial interactions and increased endothelial transcytosis (transcellular permeability)[3, 109]. Intercellular junctions (tight and adherens junctions) [78, 115] undergo post-translational modifications (e.g. phosphorylation) and disassembly from the actin cytoskeleton [116] giving rise to increased paracellular permeability. Barrier dysfunction is also accompanied by the acquisition of the cerebrovascular endothelium of a pro-inflammatory phenotype, characterized by the expression of leukocyte-endothelial adhesion molecules (e.g. E-Selectin, ICAM-1). The GTPase Rho, is emerging as a central key regulator of these cellular processes. Rho, and its effector, ROCK play a critical role in endothelial dysfunction via the regulation of the actin cytoskeleton dynamics [78, 80], caveolin-mediated endothelial transcytosis [79, 117] as well as the induction of pro-inflammatory gene expression via activation of nuclear factor κ B[118, 119]. In the current study we found that, at the molecular level, exacerbated BBB dysfunction in S1PR1 endothelial specific null mice correlated with increased levels of phosphorylated MLC, a downstream effector of the Rho-ROCK pathway implicated in blood brain barrier dysfunction [77, 78, 80]. Altogether, our data indicates that the aggravation of albumin leakage observed in endothelial specific S1PR1 null mice correlates with increased activation of Rho-ROCK pathway, implicating the role of endothelial S1PR1 signaling in the regulation of the endothelial cytoskeleton and vesicular trafficking leading to barrier dysfunction.

Other possible vascular mechanisms responsible for the worsened outcomes in endothelial specific S1pr1iECKO mice vs wild type could be differences in cerebral blood flow or in bleeding upon cerebral artery rupture. However, we did not observe differences in the CBF changes or physiological parameters during and after SAH between these two groups of mice. In addition, the amount of blood in the subarachnoid space (SAH grade), which plays an important role in the severity of brain injury and SAH outcomes was similar in both groups of mice, indicating that there were no differences in bleeding or the clearance of the subarachnoid blood. Furthermore, there were no differences in hemostasis, assessed by the tail-bleeding assay, between wild type and S1pr1iECKO mice. Thus, the exacerbation of neuronal injury upon SAH in the endothelial specific s1pr1 null mice vs wild type cannot be attributed to differences in bleeding or in cerebral blood flow regulation. Altogether our data points to exacerbation of BBB permeability (albumin leakage) as the main cellular mechanism underlying the aggravation of brain injury upon endothelial deletion of S1PR1 in the adult mice, providing proof of concept of the critical role of endothelium and blood brain barrier dysfunction in SAH outcomes.

Lastly, our study also underscores the pathophysiological relevance of the S1P pathway in humans. Using RT-qPCR in combination with mouse genetic approaches and IHC techniques previously validated in our laboratory [62], we found that S1PR1 is the most abundant S1PR transcript in the mouse brain and mouse brain microvessels and widely detected in the human cerebrovascular endothelium and brain parenchyma. Our human data, together with our mouse studies using the endothelial-specific S1PR1 null mice, highlight the therapeutic and prognostic potential of the endothelial S1P pathway in stroke. Given recent human studies which indicate that S1P-S1PR1 vasoprotective signaling [28, 30] may be limiting in cardiovascular and inflammatory diseases [17, 31–34] our present data imply that patients having limiting endothelial S1PR1 antiinflammatory signaling (e.g. with lower levels of ApoM or HDL-bound S1P), could be at risk of worsened outcomes upon stroke. In addition, our data suggests that the adverse effects reported in some subsets of multiple sclerosis patients in chronic treatment with FTY720, such as macular edema or posterior reversible encephalopathy syndrome [120–123] could be due to antagonism of endothelial S1PR1.

In summary, our study demonstrates the critical role of the endothelial S1PR1 signaling in BBB function and SAH outcomes in mice. Our human data strengthen the pathophysiological relevance of these findings and highlight the therapeutic and prognostic potential of the endothelium, more specifically the endothelial S1P signaling pathway in stroke. New strategies to modulate S1P signaling specifically in the endothelium to prevent exacerbation of BBB leakage and brain injury without compromising the immune response [17, 60, 124] could hold promise as novel neurovascular protective therapies in stroke and other pathological conditions associated with BBB dysfunction [5–13, 81–85], [86–89].

## ACKNOWLEDGEMENTS

This work was supported by internal funds provided by the Departments of Surgery and Emergency Medicine, BIDMC, the Department of Pathology and Laboratory Medicine, Weill Cornell Medicine, American Heart Association Grant-in-Aid 12GRNT12050110, NIH HL094465 and Leducq Foundation grants to TS. AI was partially supported by LeRoche foundation and the Tri-Institutional Therapeutics Discovery Institute.

We would like to thank Dan Li and Dr. Shou-ching Jaminet from the Multi-Gene Transcriptional Profiling Core Facility (Center for Vascular Biology Research, Beth Israel Deaconess Medical Center, Harvard Medical School) for the quantitative PCR analysis, Drs. Louise McCullough (University of Texas Health McGovern Medical School) and Tim Hla (Boston Children’s Hospital) for their input in the project.

The authors declare no competing financial interests.

## SUPPLEMENTARY FIGURE LEGENDS

**Supplementary Figure 1.**
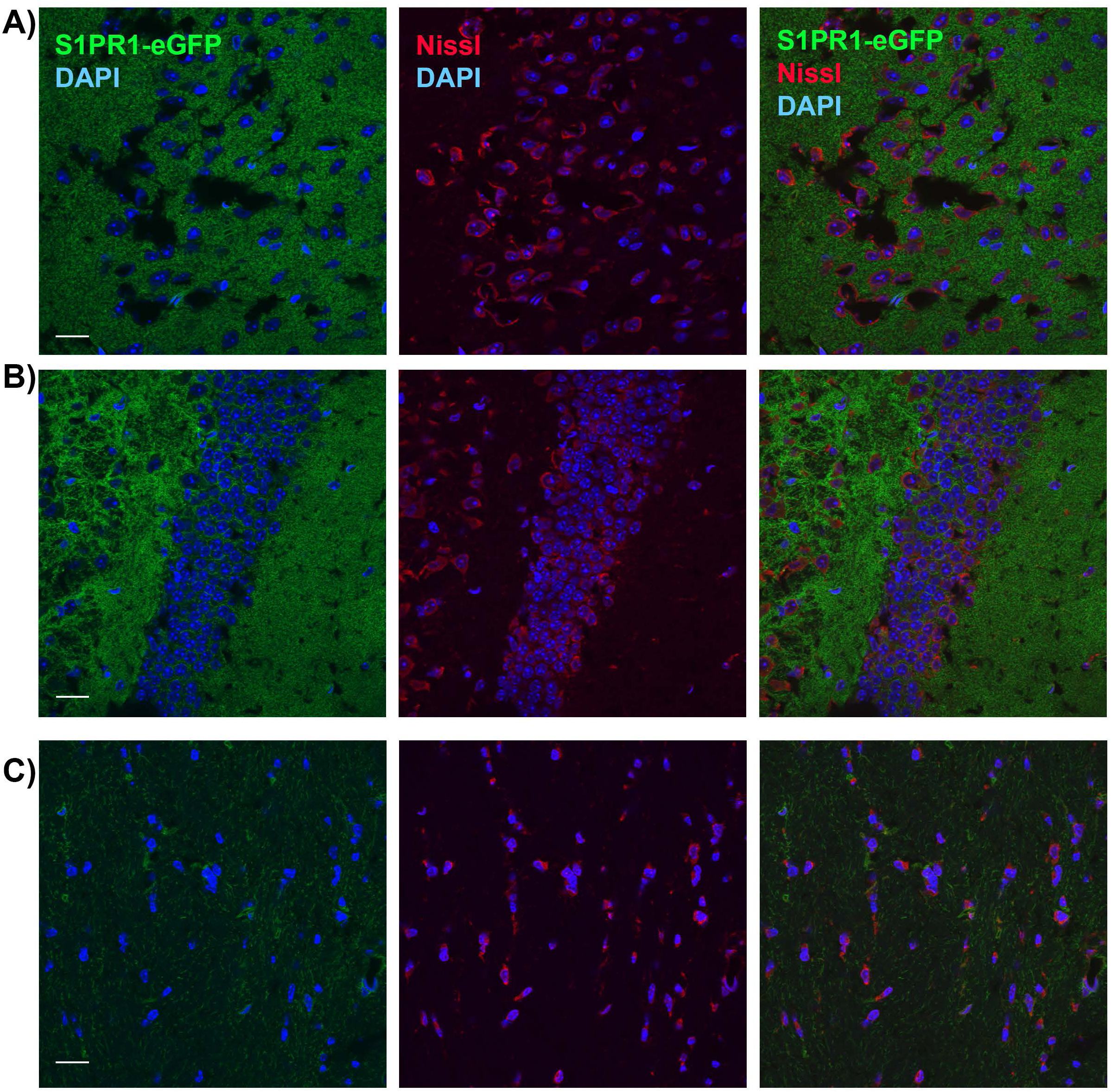
Expression of S1PR1-eGFP and Nissl stain in brain in the S1PR1-eGFP knock in mouse. S1PR1-eGFP fluorescence confocal analysis in grey matter (A, cortex and B, hippocampus) and white matter areas (C, corpus callosum) of the mouse brain. Neuronal somas are stained with NeuroTrace® 530/615 Red Fluorescent Nissl (red channel). Note that S1PR1-eGFP is expressed in neurons and localized around the soma, mainly in the neuropil (A, B) and axons (C). Sections were imaged by using an FluoView FV10i confocal microscope (Olympus, Japan), (original magnification, × 60). Scale bar 20μm. Representative pictures are shown. N=5-6

**Supplementary Figure 2.**
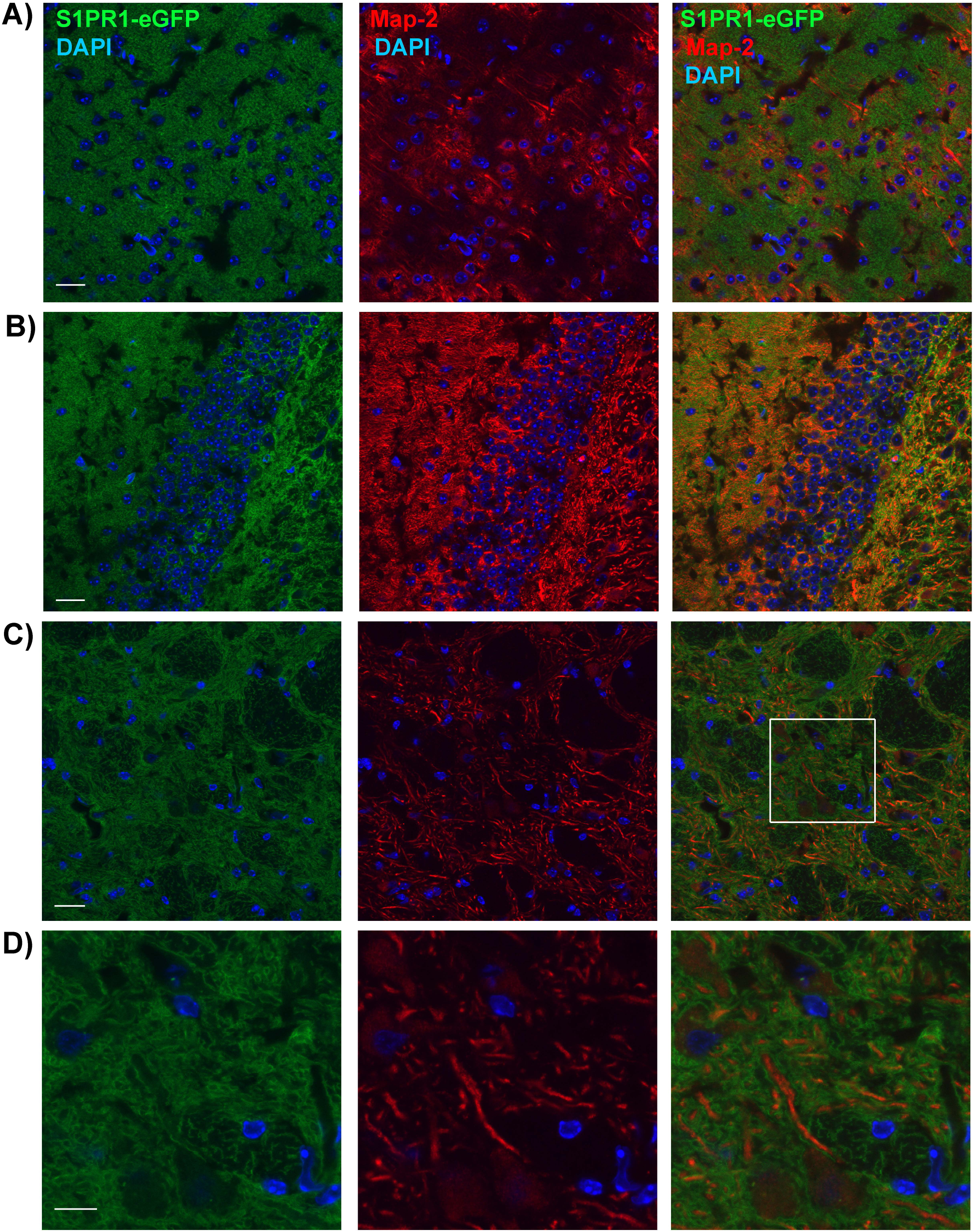
Expression of S1PR1-eGFP and MAP-2 immunofluorescence in neurons in the S1PR1-eGFP knock in mice. S1PR1-eGFP fluorescence confocal analysis in grey matter (A, cortex and B, hippocampus) and white matter areas (C, D, internal capsule) of the mouse brain. Notice that S1PR1-eGFP signal is localized in the dendrites and neuronal processes (MAP-2 positive). D) Inset shown in panel C (digital zoom). Image shows localization of S1PR1-eGFP around the microtubules. Sections were imaged by using an FluoView FV10i confocal microscope (Olympus, Japan), (original magnification, × 60). Scale bar 20μm. Representative pictures are shown. N=5-6

**Supplementary Figure 3.**
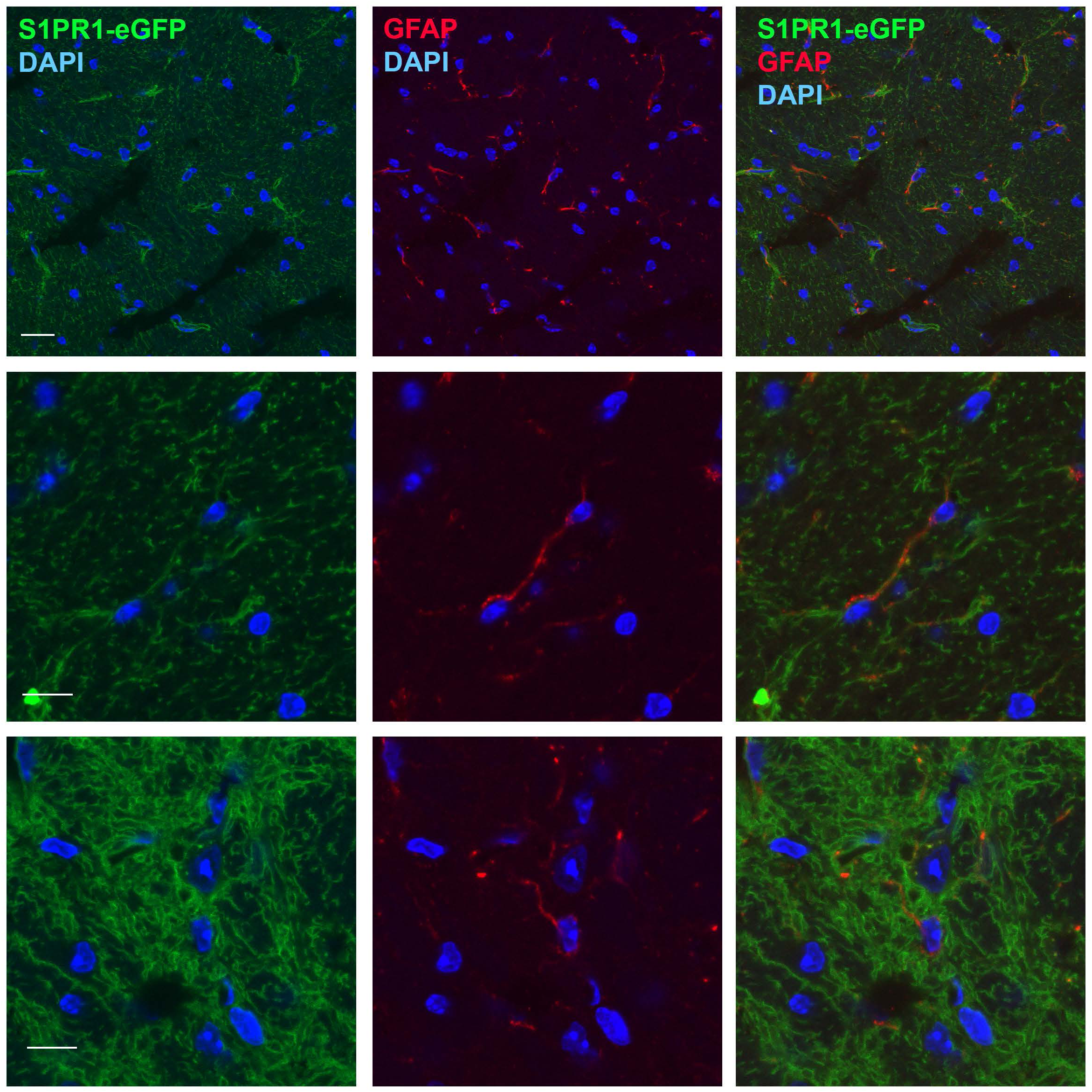
Expression of S1PR1-eGFP and GFAP immunofluorescence in astrocytes in the S1PR1-eGFP knock in mice. S1PR1-eGFP fluorescence confocal analysis in white matter areas (A, B, corpus callosum, C, internal capsule) of the mouse brain. Arrows indicate fibrous astrocytes positive for S1PR1-eGFP. B) Digital zoomed image of an astrocyte positive for S1PR1-eGFP. C) Digital zoomed image of an astrocyte negative for S1PR1-eGFP.Sections were imaged by using an FluoView FV10i confocal microscope and a 60x objective (Olympus, Japan) Scale bar 20μm. Representative pictures are shown. N=5-6

**Supplementary Figure 4.**
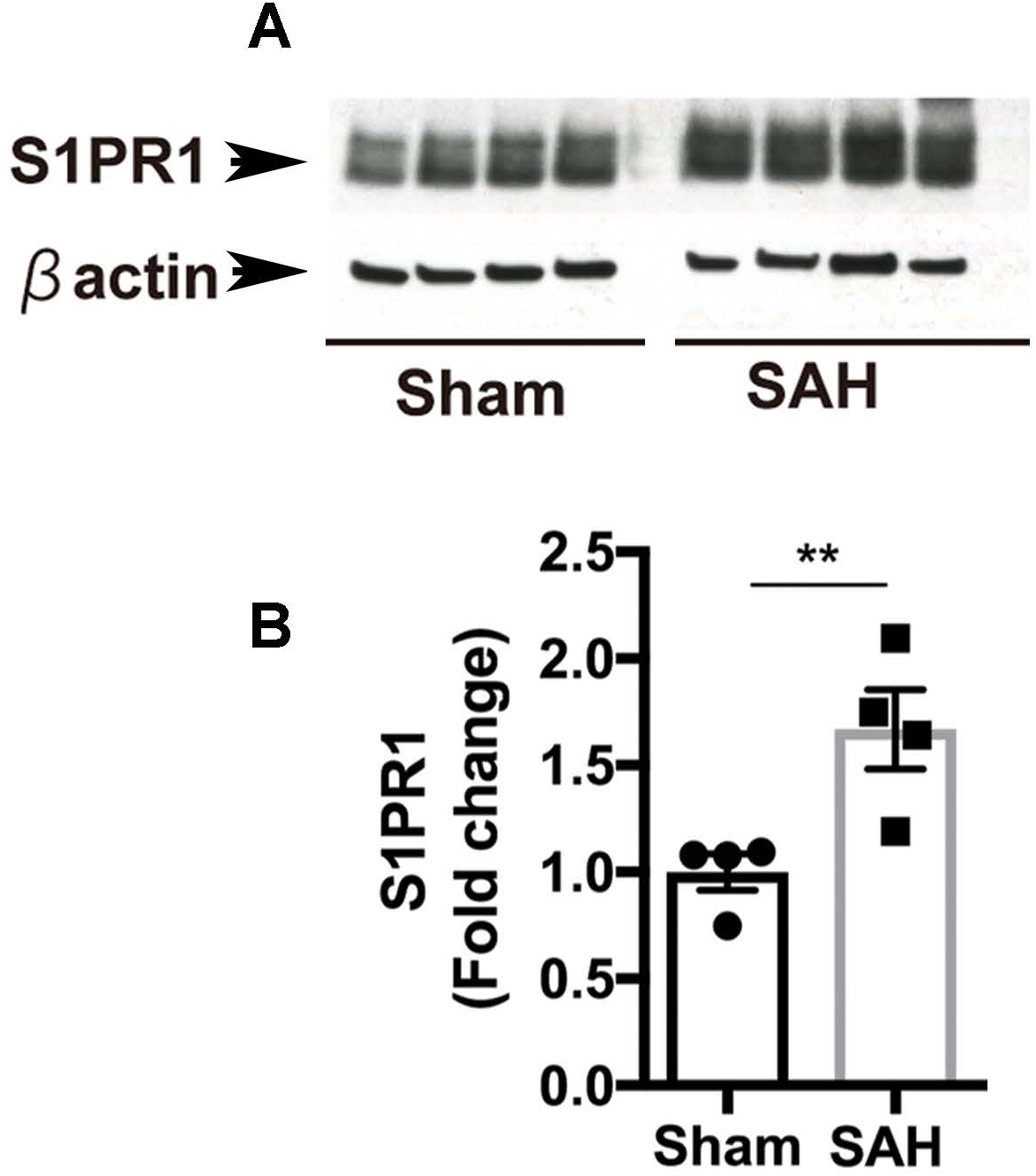
S1PR1 protein levels in cortical microvessels are increased 24h after SAH. Western blot analysis for S1PR1 in brain microvessels. 24 after subarachnoid hemorrhage (SAH) cortical microvessels were isolated and S1PR1 levels were detected by western blot (*n* = 4). Immunoblot image (A) and quantification of S1PR1 (B) are shown. βactin bands are used as a loading control and for normalization in quantification. Individual values and mean ±SEM are shown. \*\**P* < 0.01, t test.

